# Pleiotropic effects on *E. coli* physiology of the AraC-like regulator from prophage origin, AppY

**DOI:** 10.1101/2022.02.28.482378

**Authors:** Naoual Derdouri, Nicolas Ginet, Yann Denis, Mireille Ansaldi, Aurélia Battesti

## Abstract

Bacterial genome diversity is largely due to prophages, which are viral genomes integrated into the genomes of bacteria. Most prophage genes are silent, but those that are expressed can provide unexpected properties to their host. Using as a model *E. coli* K-12, that carries 9 defective prophages in its genome, we aimed at highlighting the impact of viral genes on host physiology. We focused our work on AppY, a transcriptional regulator encoded on the DLP12 prophage. By performing RNA-Seq experiments, we showed that AppY production modulates the expression of more than 200 genes; among them, 13 were identified by ChIP-Seq as direct AppY targets. AppY directly and positively regulates several genes involved in the acid stress response including the master regulator gene *gadE*, but also *nhaR* and *gadY*, two genes involved in biofilm formation. Moreover, AppY indirectly and negatively impacts bacterial motility by favouring the degradation of FlhDC, the master regulator of the flagella biosynthesis. As a consequence of these regulatory effects, AppY increased acid stress resistance and biofilm formation while also causing a strong defect in motility. We therefore demonstrate here that AppY is a central regulator from phage origin that controls the expression of bacterial master regulators to provide benefits to *E. coli* under stress conditions. Our research shed light on the importance to consider the genetic dialogue occurring between prophages and bacteria to fully understand bacterial physiology.

## INTRODUCTION

Metagenomic studies have revealed that viruses infecting bacteria, called bacteriophages or phages, are the most abundant and diverse biological entities on the planet (1,2). They are found in various environments such as ocean, soil or air and more recently, they have been shown to be ubiquitously present in the human body as the major constituents of the “virome” (2–4). Phages can be classified according to their life cycles (1). Virulent phages only perform a lytic cycle: after host recognition and genome injection, they hijack the host machineries to produce new virions, leading to host lysis when released. Temperate phages can adopt a lytic cycle or alternatively and, depending on the host physiology, a lysogenic cycle. In the lysogenic cycle, the viral genetic material can be integrated into the host chromosome as a prophage and be vertically transferred to the host progeny (5). Once integrated, prophage encoded genes can be silenced by viral or bacterial regulators to avoid detrimental effects on host physiology. During evolution, some of these prophage genes are lost due to mutation acquisition, deletion and recombination events (6). As a consequence, some prophages become defective, *i. e.* they lose their ability to resume a lytic cycle. However, these defective prophages should not only be considered as genetic material in decay since they still contain intact genes that retain some functions and can be expressed under specific conditions. Some of these genes, called morons, do not participate in the phage cycle but provide benefits to their host (7). Although the contribution of these morons to bacterial pathogenicity has been studied, considerably less data are available to explain their impact on bacterial fitness or stress resistance (8–10). However, having a full picture of functional interactions linking prophages to their host is essential to fully understand bacterial physiology and assess the contribution of horizontally transferred functions.

Transcriptional regulators drive gene expression and consequently can have a major impact on bacterial physiology. Out of the roughly 300 transcriptional regulators encoded in the *E. coli*strain MG1655 genome, only 185 have been experimentally characterized (11–13). This strain contains 9 defective prophages in which 9 transcriptional regulators unrelated to classical regulators of phage cycle such as C1 or Cro have been identified (14). Although the regulon of each of these prophage-encoded regulators has not been characterized in detail, data obtained so far suggest that a majority of them regulates the expression of prophage genes (14). One notable exception is the transcriptional regulator AppY.

*appY* is located in the defective lambdoid prophage DLP12 integrated into the *argU* tRNA gene (15,16). DLP12 is the most prevalent prophage in *E. coli* strains and is also found in *Shigella* genomes (17). *appY* expression is induced under anaerobiosis, phosphate and carbon starvations as well as during entry into stationary phase. Depending on the environmental conditions, its expression relies on the global regulators ArcA, H-NS, RpoS or the two-component system DpiA/B (18–21). AppY is a transcriptional regulator from the AraC/XylS family, which is one of the most represented family of transcriptional regulators in bacteria (22). Regulators from this family contain a characteristic DNA binding domain composed of two helix-turn-helix motifs. Most of them also contain a second domain involved in dimerization and/or effector binding. Members of this family are usually involved in general metabolism, virulence or stress responses (22–24). To date, the exact role of AppY in cell physiology remains elusive and only a few AppY targets have been identified. Indeed, AppY has been shown to induce the expression of two operons located on the host chromosome: the *hya* operon coding for the hydrogenase 1 and the *app* operon coding for the cytochrome bd-II oxidase (15,18,19,25–28). More recently, it has been shown that AppY overproduction leads to the stabilization of RpoS, the main sigma factor in stationary phase and the master regulator of the general stress response by a mechanism that has not been elucidated thus far (29,30). Finally, a qualitative study has shown that AppY overproduction affects positively or negatively the level of more than 30 proteins in *E. coli* although none of these proteins were identified in that work (15).

Here, we aim to determine the contribution of AppY, a regulator acquired by horizontal gene transfer, to host physiology. By using global approaches (RNA-Seq and ChIP-Seq) as well as forward genetics, we identified genes directly and indirectly regulated by AppY. We showed that AppY contributes to bacterial survival under low pH conditions, biofilm formation and repression of bacterial motility. We identified and characterized the regulatory pathways leading to each of these adaptive responses triggered by AppY. Therefore, our study provides molecular insights into AppY integration into the *E. coli* regulatory network. It also highlights how a prophage-encoded transcriptional regulator integrates in a complex manner in to the host regulatory network and how it benefits its host, allowing it to cope with changing environmental conditions.

## RESULTS

### Global picture of the AppY regulon

As mentioned above, AppY overproduction stabilizes the master regulator RpoS, which itself regulates more than 500 genes in *E. coli* (30,31). Therefore, to define the AppY regulon and avoid the identification of genes under RpoS control, we overproduced AppY from an inducible plasmid in a strain deleted for *rpoS.* Variations in RNA expression were measured by RNA-Seq experiments. Using this approach, we identified more than 200 genes whose expression was significantly (5-Fold) modulated upon AppY overproduction (Supplementary Table 1). 65 genes, whose expression varied by 10-fold or more when AppY was overproduced, were classified based on their biological functions according to Ecocyc (Figure 1A). Up-regulated genes fell into 5 distinct functional groups: metabolism, regulation, transport, respiration, and stress adaptation (Figure 1A). We observed a massive induction of the two known AppY targets involved in respiration, the *hya* and *app* operons (containing 6 and 4 genes respectively), therefore validating our strategy to identify the AppY regulon (Figure 1B, blue bars) (32). Among the 21 genes involved in stress adaptation, 19 genes are involved in acid stress resistance; 17 genes out of the 19 genes code for proteins belonging to the glutamate-dependent acid resistance system 2 (AR2), the major acid resistance pathway in *E. coli* (33) (Figure 1B, pink bars). Down-regulated genes were also identified in these experiments and divided into two categories: metabolism and motility. All of the 4 down-regulated metabolism genes were part of diverse metabolic pathways with no obvious link between them while the 13 genes linked to motility all code for proteins involved in flagella synthesis. In addition, 11 other genes involved in flagella synthesis were also down-regulated between 3 to 10 times (Figures 1B, light green bars). Overall, these data suggest two unexpected roles for AppY: it contributes to bacterial adaptation under low pH by inducing the glutamate-dependent response and it negatively regulates bacterial motility by down-regulating flagellar genes.

**Figure 1 :**
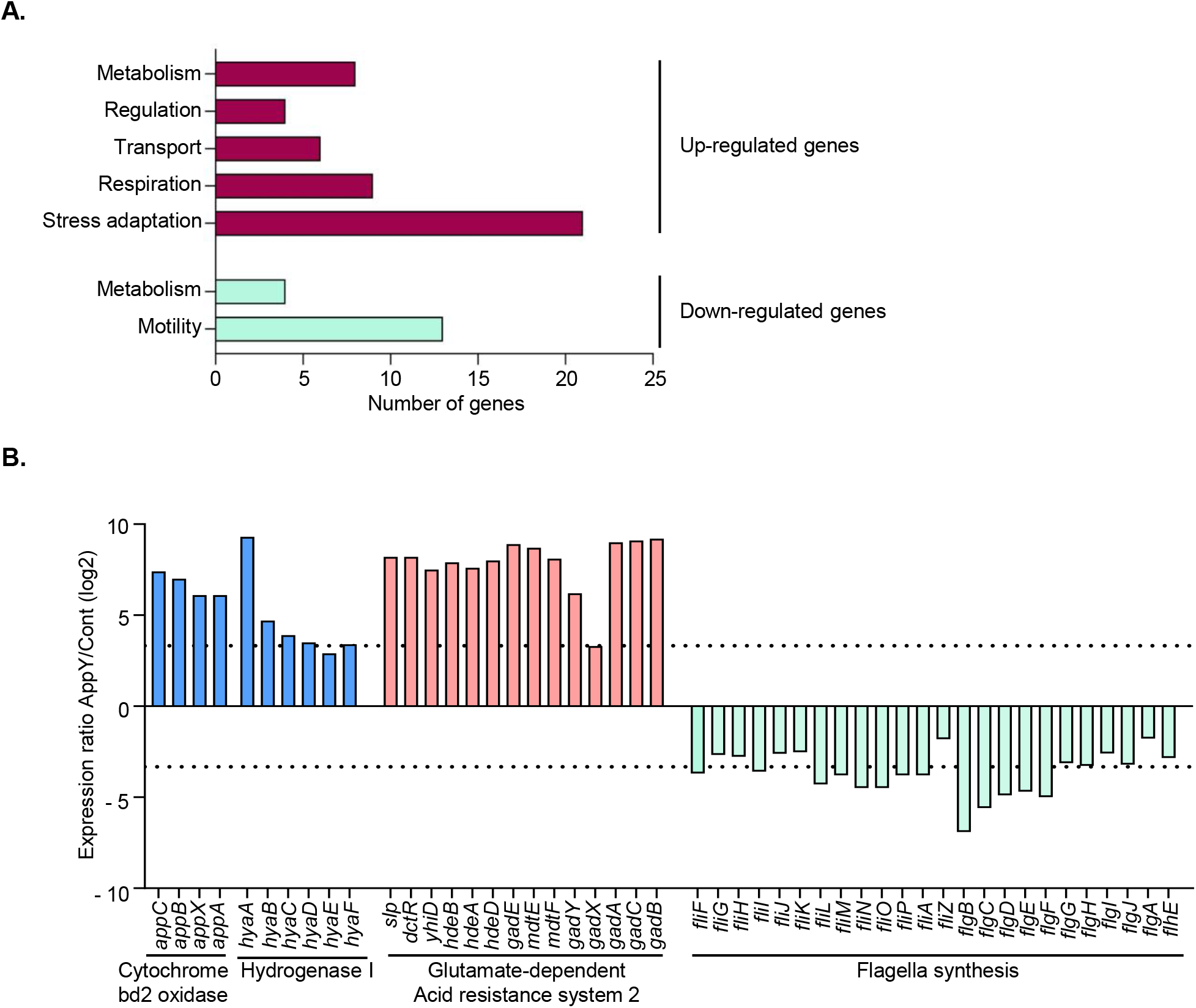
Identification of AppY regulon by RNA-Seq experiments. MG1655 *rpoS::tet* strain transformed with plasmids pQE80L or *pQE80L-appY* were grown to an OD_600_ ~0.6 and *appY* expression was induced with 1 mM IPTG for 1 hour. RNA were purified, reverse-transcribed and sequenced. The data shown are representative of three independent experiments. A. Global overview of the function of genes regulated by AppY. Genes up-regulated (dark red) and down-regulated (light green) at least 10 times when AppY was overproduced were classified according to their biological functions using Ecocyc. B. AppY mainly regulates genes involved in respiration, acid resistance and flagellar synthesis. Genes in blue are involved in aerobic respiration and are known AppY targets, genes in pink belong to the AR2 system and genes in green are involved in flagella synthesis. The dashed lines indicate 10-fold induction or repression.

### Identification of genes directly regulated by AppY

RNA-Seq experiments gave a global overview of the physiological changes occurring in the cell when AppY was overproduced for an hour but did not provide information on AppY direct targets. To identify the genes that are directly regulated by AppY, ChIP-Seq experiments were performed. Briefly, the AppY protein with a 3Flag-tag at its C-terminal end was produced from an inducible plasmid in the *rpoS::tet* strain. Note that the 3Flag-tag did not affect AppY functionality (Figure S1A). After stabilization of the AppY-3Flag/DNA complexes using formaldehyde, chromosomal DNA was fragmented and cells lysed by sonication. AppY-3Flag/DNA complexes were then recovered by affinity purification using the 3Flag-tag, and bound DNA was sequenced. To get rid of false positive results, the experiment was performed in parallel with an AppY-3Flag mutant unable to bind DNA. Such mutants affected in their DNA binding function have been well-characterized in other AraC-transcriptional regulators (34). Based on sequence alignments, we identified a lysine residue at position 170 as potentially involved in AppY DNA-binding (Figure S1B). We used site-directed mutagenesis to generate the AppY_K170E_ mutant and checked that its production was similar to that of the wild-type protein (Figure S1C). We then confirmed that the K170 residue was critical for AppY function, since the AppY_K170E_ did not allow the expression of the two transcriptional fusions used as positive controls *P_appC_-gfp* and *P_hyaA_-gfp* (Figure S1A).

Using ChIP-Seq, 13 AppY binding-sites were identified (Figure 2A). The detection of such sites in the promoter regions of *hyaA* and *appC* was in accordance with the RNA-Seq data and demonstrated for the first time, to our knowledge, that AppY directly regulated these operons (Figure 2B). Strikingly, no AppY binding site was identified in the promoter region of the genes involved in flagella synthesis that were heavily repressed in RNA-Seq experiments (Figure 1B), suggesting an indirect regulation of these genes by AppY (Figure 2A). In contrast, AppY was clearly able to bind upstream of six genes that are involved in acid stress resistance (*slp*, *gadE*, *gadY*, *adiC*, *glsA* and *yiiS*). Among them, 3 genes are located in the acid fitness island and involved in the AR2 pathway: *slp*, *gadY* and *gadE* (Figures 2A and C). GadE is the master regulator of the AR2 pathway, suggesting that its gene expression regulation by AppY can have broad consequences on the AR2-dependent acid response (35–37). In addition to its role in acid stress, GadY is a non-coding RNA also involved in biofilm formation (38–40). We identified another AppY target, *nhaR*, also involved in biofilm synthesis, suggesting a regulatory role of AppY in this pathway (41). Finally, for the 5 remaining genes that we identified by the ChIP-Seq approach (*groSL*, *acrZ*, *prc*, *uspD* and *rrsG*), we did not find any obvious physiological links with the other targets; therefore, additional experiments are needed to confirm these targets and to understand the consequences of their induction when AppY is overproduced.

**Figure 2 :**
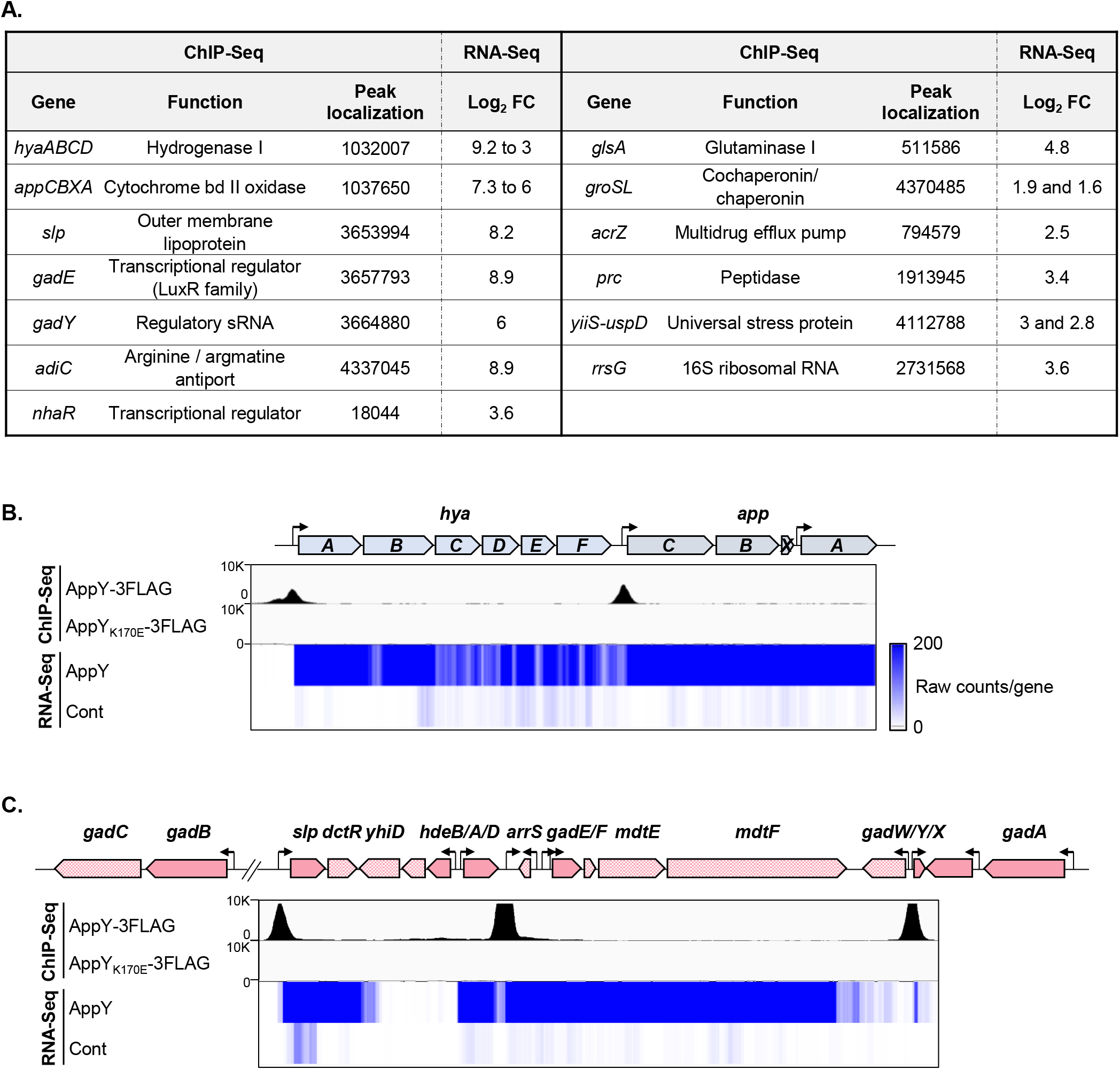
Identification of AppY binding sites by ChIP-Seq experiments. A. AppY direct targets. Name, function, peak localization of AppY binding on *E. coli* chromosome and the log_2_ fold change (FC) obtained in RNA-Seq experiments are summarized. B. Regulation of the *hya* and *app* operons. MG1655 *rpoS::tet* strain containing the plasmids pQE80L, pQE80L-*appY*-3FLAG or *appY*_K170E_-3FLAG were grown at 37°C to an OD_600_~0.2 and *appY* expression was induced with 0.05 mM IPTG for 1 hour. The genetic loci, the ChIP-Seq data obtained with AppY-3FLAG and AppY_K170E_-3FLAG and the gene expression profiles obtained by RNA-Seq with AppY or the vector control are shown from the top to the bottom. The black peaks observed in ChIP-Seq experiments correspond to AppY binding to the DNA; the intensity of the blue color in RNA-Seq panels represents the number of raw counts per gene. The data are representative of three independent experiments. C. Regulation of the acid fitness island. Experiments were performed and presented as described in B.

Overall, the Chip-Seq results combined with those of the RNA-Seq suggest that AppY is involved in acid stress adaptation, biofilm formation and motility inhibition. Hence, we explored the role of AppY in each of these pathways.

### AppY directly and indirectly induces the expression of genes from the AR2 system

The identification of *gadE* as a direct AppY target suggests that the induction of several genes from the AR2 pathway observed in the RNA-Seq experiment was probably GadE-dependent. To test this hypothesis, we used *gfp*-transcriptional fusions in strains deleted of *rpoS* only or both *rpoS* and *gadE* (Figure 3A). For genes organized in an operon, we used fusions containing the promoter region in front of the first gene of the operon (Figure 2C, deep pink genes in the genetic loci). Background levels of fluorescence were measured in strains containing the control plasmid (no promoter upstream of *gfp*) (Figure 3A, “cont” bars), or containing the fusions of interest in the absence of AppY overproduction (Figure 3A, hatched bars). The *appC* fusion was used as a positive control since we observed a direct AppY binding in its promoter region (Figures 2B and 3A). When AppY was overproduced in the strain deleted of *rpoS*, fluorescence levels were considerably increased for most of the fusions (from 4 to 34-fold; Figure 3A, grey bars). These results are consistent with the RNA-Seq data showing induction of these genes in the presence of AppY (Figure 1B). Only a mild effect was seen for *hdeA* and *gadX* (1.6 and 1.3-fold increase, respectively). One possible explanation is that the increased level of *hdeA* and *gadX* mRNA measured with the RNA-Seq experiment in the presence of AppY is not due to an increase in transcription but to a post-transcriptional regulation. For *gadX*, this regulation could be dependent on the small RNA *gadY* that is known to stabilize *gadX* mRNA and that we identified as a direct AppY target (38) (Figure 2A and 2C). In the strain deleted of *rpoS* and *gadE*, AppY overproduction led to the induction of *appC, slp, gadE* and *gadY*, the direct AppY targets we identified by ChIP-Seq (Figure 3A). However, AppY overproduction failed to induce the expression of *hdeD*, *gadB* and *gadA* in this genetic background. These results confirm that AppY directly induces *gadE* expression which in turn activates its own regulon including *hdeD, gadB* and *gadA*. Altogether, these experiments identify AppY as a new player leading both directly and indirectly to the activation of the AR2 pathway.

**Figure 3 :**
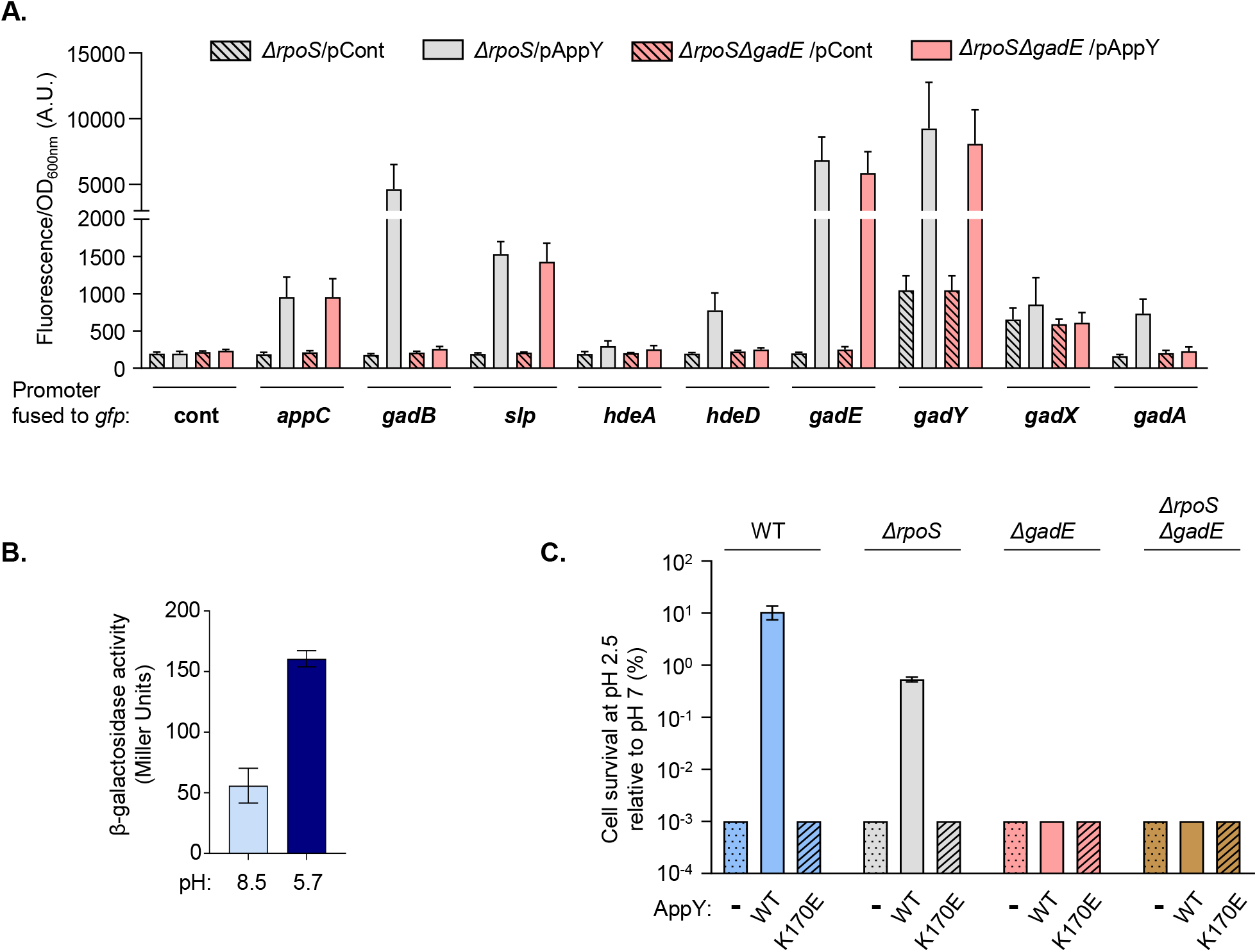
AppY contributes to acid stress response. A. *gadE*-dependent AppY targets. MG1655 strains deleted of *rpoS* (Δ*rpoS*,grey) or both *rpoS* and *gadE (ΔrpoSΔgadE*,pink) and containing the pQE80L empty vector (pCont, hatched) or *pQE80L-appY* (pAppY, plain) were transformed with the indicated transcriptional *gfp* fusions (94). Relative fluorescence intensities were measured after overnight growth at 37°C in LB plus 0.05 mM IPTG to induce *appY* expression. Data are means +/− standard deviation (n=3). A.U., arbitrary units B. Induction of *appY* expression under acid stress conditions. An MG1655 strain carrying a chromosomal *appY-lacZ* translational fusion was cultured overnight in LBK pH=7, diluted 1:1000 into LB at pH 8.5 (light blue) or 5.7 (dark blue) and incubated at 37 °C under aerobic conditions until OD_600_ ~0.4. The activity of the *appY* fusion was determined using the Miller assay (90). The mean from three replicates is presented; the error bars represent the standard deviation (SD). C. AppY overproduction confers resistance to acid stress. WT (blue), Δ*rpoS* (grey), Δ*gadE*(pink) or Δ*gadEΔrpoS* (brown) strains transformed with pQE80L (dotted), pQE-*appY*_WT_ (plain) or pQE-*appY*_K170E_(hatched) were grown to OD_600_=1 in LB (pH 7.0) with 1 mM IPTG. Cells were diluted 1:200 into LB (pH 2.5) and incubated for 1 h at 37 °C. Cells were spotted on plates to evaluate the number of cells that survive acid stress compared to the initial number of cells. Data are means +/− standard deviation (n=3).

### *appY* induction contributes to *E. coli* survival in acidic environment

Since AppY directly regulates *gadE* leading to the activation of the AR2 pathway, we could expect that *appY* itself is expressed under acidic conditions. Looking at the literature, we found two global studies reporting an induction of *appY* expression during a shift from pH=8.5 to pH=5.7; *appY* expression was further increased in the absence of oxygen (42,43). Using a strain containing a chromosomal *appY-lacZ* translational fusion, we followed *appY* expression through beta-galactosidase assays under the growth conditions used in these studies. We measured a 2.8-fold increase in the activity of the fusion when the strain was grown at pH=5.7 compared to pH=8.5, validating the fact that *appY* expression was induced in acidic environment (Figure 3B). However, we did not detect a higher induction in the absence of oxygen (Figure S2A). The two-component system EvgA/EvgS has been shown to be involved in the sensing of acidic environment and activation of the AR2 system (44). To determine if EvgA/S and AppY belonged to the same regulatory circuit, we measured the activity of the *appY-lacZ* fusion at pH=5.7 or 8.5 in a strain deleted of *evgA* (Figure S2B). The absence of *evgA* did not affect *appY* induction at pH=5.7, leading to the conclusion that *appY* is part of a regulatory pathway independent of the two-component system EvgA/S.

We then assessed the contribution of AppY to acid resistance under conditions known to activate the AR2 system. This system requires glutamate to function: in the absence of glutamate, the AR2 system is not active and the cell cannot survive at pH=2.2 (35,45,46). As a control, we used a strain deleted of *gadC* that codes for the transporter allowing glutamate entry in the cells. As expected, a strain deleted of *gadC* did not grow with or without glutamate (Figure S2C). The WT and the Δ*appY* strains were unable to grow in the absence of glutamate but grew when glutamate was added without any significant difference (Figure S2C), indicating that in this experimental setup AppY is not essential for AR2 system activation. We then postulated that if AppY was indeed involved in acid stress management, the benefit of AppY overproduction for the host could be an increased cell survival at low pH. To test this hypothesis, AppY wild-type or the K170E mutant were overproduced in the wild-type MG1655 strain; the cultures were then shifted from pH=7 to pH=2.5 for one hour and cell survival assayed by plating. Acid stress severely affected survival of the wild-type strain containing an empty vector after one hour at pH=2.5 (Figures 3C, blue bars and S2D). Strikingly, AppY overproduction massively increased cell survival by at least 100-fold under the same conditions. The increase in acid stress resistance was not observed with the AppYK170E mutant that does not bind DNA (Figures 3C (hatched blue bar) and S2D). These results confirm that AppY confers acid resistance and that this effect depends on its transcriptional regulator function. The effect of AppY overproduction was also tested in a Δ*rpoS* genetic background. In this context, we found that AppY overproduction still increased cell survival but 10-fold less than in a WT background (Fig. 3C, compare WT (blue bars) and Δ*rpoS* (grey bars)). This difference can be attributed to a higher sensitivity to acid stress of the strain lacking *rpoS*. To definitively link the positive effect of AppY on survival to the induction of the AR2 system, we repeated the same experiments in strains deleted of *gadE*. As expected, AppY overproduction did not increase bacterial survival in this genetic context (Figures 3C (pink and brown bars) and S2D). Overall, the data presented here demonstrate that AppY is involved in *E. coli* survival to acid stress, mainly by inducing *gadE* expression, hence the AR2 system.

### AppY overproduction increases biofilm formation *via* the simultaneous induction of *gadY* and *nhaR*

Out of the 13 direct AppY targets identified by ChIP-Seq, *gadY* and *nhaR* have been shown to be involved in biofilm formation through the up-regulation of the *pgaABCD* operon, responsible for the synthesis of the adhesin poly-ß-1,6-N-acetyl-D-glucosamine (40,41). In order to test if AppY overproduction leads to biofilm formation, MG1655 *rpoS::tet* strain was transformed either with the pQE80L empty vector or with the same vector containing *appY* WT or the mutant affected in DNA-binding. Cells were grown for 24 hours at 30°C and biofilm formation was quantified with crystal violet staining and OD_550_ measurement. This quantification clearly showed that AppY overproduction led to a 3-fold increase in biofilm formation (Figures 4A, grey bars and S3A). AppY_K170E_ overproduction had no effect on biofilm showing that the observed phenotype is linked to the regulatory function of AppY (Figures 4A, grey hatched bars and S3A). To confirm that AppY acted on biofilm through GadY and NhaR, we deleted the corresponding genes in the MG1655 Δ*rpoS* background. AppY overproduction in Δ*rpoS* Δ*gadY* or in Δ*rpoS* Δ*nhaR* strains, did not lead to biofilm formation suggesting that GadY and NhaR are both essential in this pathway (Figures 5A pink and purple bars respectively and S3A). In order to quantify the contribution of these two regulators for biofilm formation, we cloned *gadY* and *nhaR* under their own promoter and co-transformed them with either pQE80L, pQE80L-*appY*_WT_ or pQE80L-*appY*_K170E_ in strains deleted of *gadY* or *nhaR*. In the presence of AppY, *nhaR* and *gadY* should be expressed from the plasmids and complement the biofilm formation defect of these two strains. In the MG1655 Δ*rpoS* Δ*gadY* strain, the plasmid expressing *gadY* showed no effect on biofilm when co-transformed with the control vector (Figure 4B, pink dotted bar and S3B). However, the production of AppY led to the expression of *gadY* and restored biofilm formation (Figure 4B, plain pink bar and S3B). As expected, no biofilm was observed with AppY_K170E_. Interestingly, in this genetic background, we did not observe an increase in biofilm with the plasmid expressing *nhaR*, even in the presence of AppY (Figure 4B, purple bars and S3B). Altogether, this first set of experiments shows that NhaR alone is not sufficient to induce biofilm formation and that AppY contributes to biofilm formation through *gadY* induction. We then performed the same experiments in a MG1655 Δ*rpoS* Δ*nhaR* strain. In this background, GadY alone was not sufficient to induce biofilm formation, even with AppY (Figure 4C, pink bars and S3C). In the presence of *nhaR* and the control vector we observed a slight increase in biofilm formation; the same result was obtained with AppY_K170E_ (Figure 4C and S3C). However, biofilm was fully restored only when both AppY and NhaR were produced at the same time (Figure 4C, plain purple bar). Overall, these data confirm that AppY favors biofilm formation by activating simultaneously the expression of the transcriptional regulator gene *nhaR* and the small RNA GadY.

**Figure 4:**
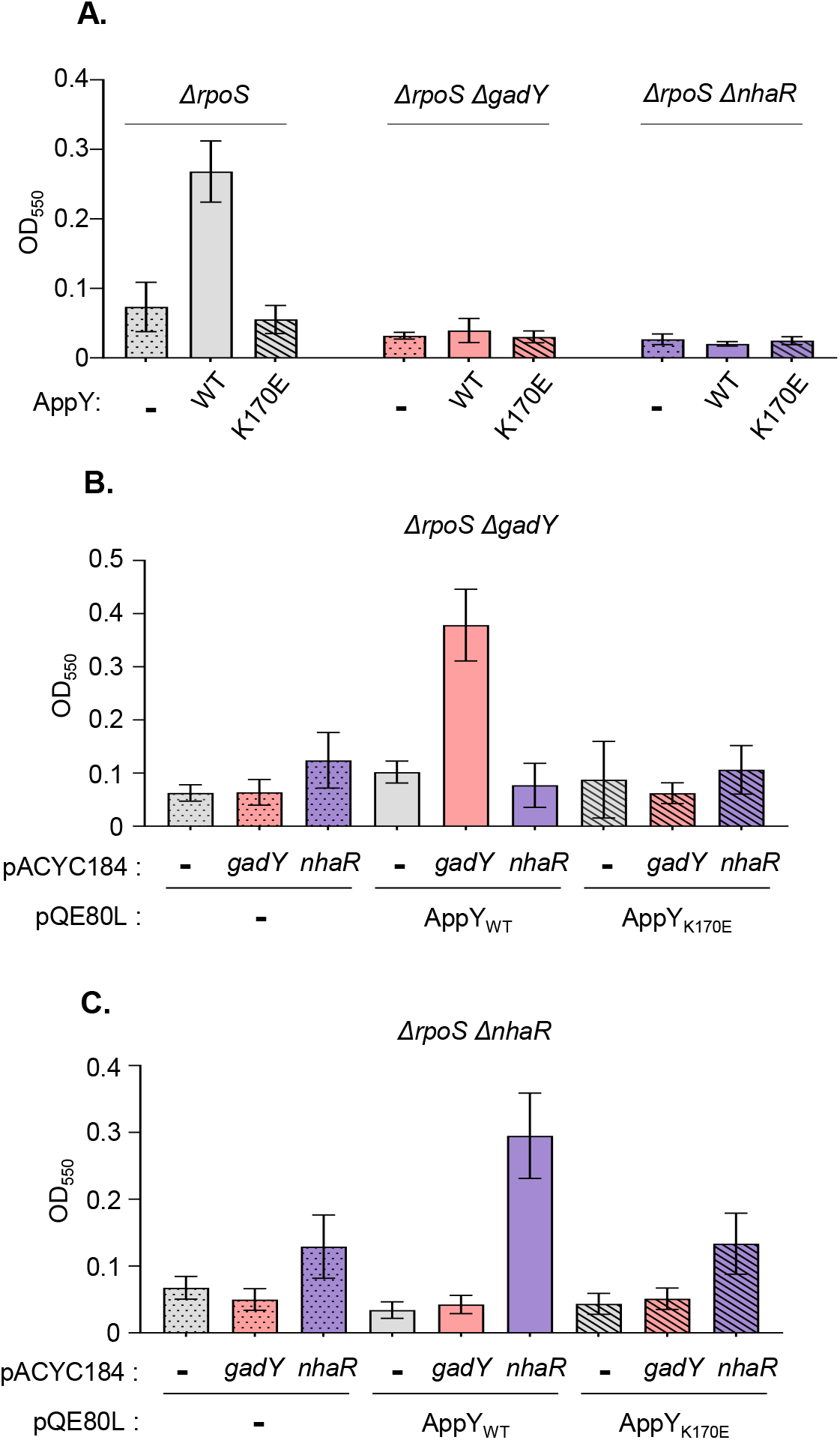
AppY favors biofilm formation through *nhaR* and *gadY*induction. The indicated strains were grown in LB with 0.5 mM IPTG at 30°C without shaking for 24 hours. Biofilm formation was visualized using crystal violet staining and quantified by measuring the optical density at 550 nm (OD_550_). The mean from three replicates is presented; the error bars represent the standard deviation (SD). A. Biofilm formation dependent on AppY, GadY and NhaR. MG1655 Δ*rpoS* (grey), MG1655 Δ*rpoS* Δ*gadY*(pink) or MG1655 Δ*rpoS ΔnhaR* (purple) were transformed with the pQE80L empty vector (-, dotted) or containing *appY* (WT, plain) or mutant (K170E, hatched). B. Complementation of MG1655 Δ*rpoS ΔgadY* with *gadY and nhaR* expressed under their own promoters. MG1655 Δ*rpoS ΔgadY* strain was co-transformed with the following plasmids: pQE80L (-, dotted), containing *appY* (WT, plain) or mutant (K170E, hatched) and a pACYC184 construct (empty vector, −, grey), containing *gadY* (pink) or *nhaR* (purple). C. Complementation of MG1655 Δ*rpoS ΔnhaR* with *gadY and nhaR* expressed under their own promoters. MG1655 Δ*rpoS ΔnhaR* strain was co-transformed with a pQE80L construct (empty vector (-, dotted line), containing *appY* (WT, plain) or mutant (K170E, hatched)) and a pACYC184 construct (empty vector (-, grey), containing *gadY* (pink) or *nhaR*(purple)).

### AppY promotes the degradation of the master regulator FlhC, leading to a strong defect in bacterial motility

According to the RNA-Seq experiments, AppY overproduction strongly down-regulates several genes involved in flagellar formation and motility (Figure 1). However, the ChIP-Seq data showed no AppY binding site upstream of these genes suggesting that this regulation was indirect (Figure 2A). To confirm the negative effect of AppY on the expression of flagellar genes, we quantified the level of expression of the first gene of each operon when AppY was overproduced using qRT-PCR (Figure 5A; plain arrows). *appC* was used as a positive control since we have demonstrated that it is directly and positively regulated by AppY (Figures 2B and 5A). In agreement with the RNA-Seq data, expression of all of the *fli* and *flg* genes tested was reduced between 2.5 and 16-fold in the presence of AppY. This decrease in gene expression was not observed with AppY_K170E_ confirming that an intact DNA binding domain is needed to exert this repression (Figure 5A). We then assayed the consequences of this repression on *E. coli* motility. We transformed pQE80L-*appY*, pQE80L-*appY*_K170E_ or the control vector pQE80L into the MG1655 Δ*rpoS* strain and performed a classical swimming test on soft-agar plates. In the absence of inducer, all strains were equally motile (Figure 5B, light green bars). In contrast, in the presence of IPTG, AppY overproduction inhibited cell motility (Figure 5B, dark green bars). Here again, motility inhibition was not observed with the AppY _K170E_ mutant (Figure 5B).

**Figure 5:**
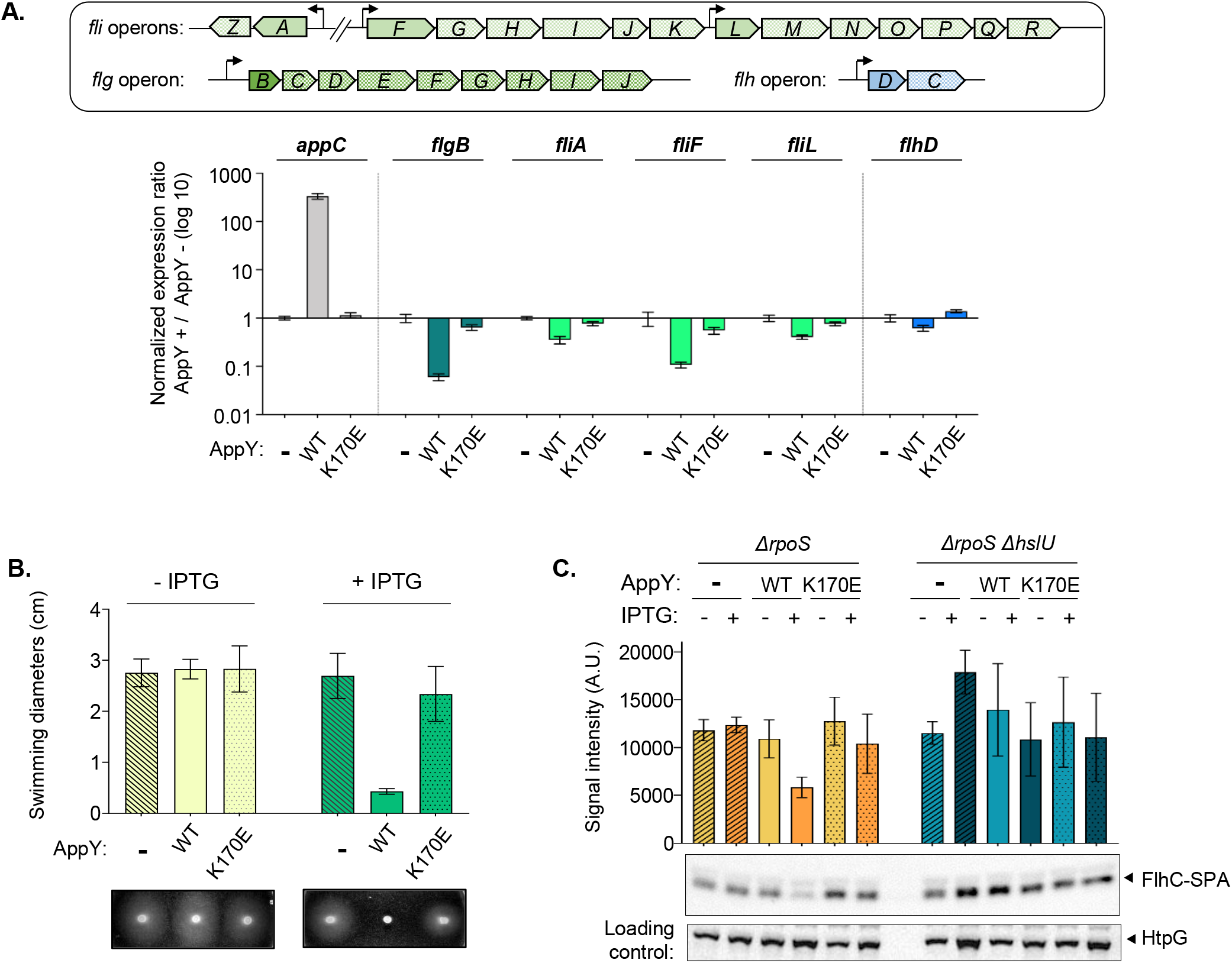
AppY indirectly reduces bacterial motility. A. Expression of flagellar genes is down-regulated by AppY overproduction. MG1655 Δ*rpoS* strain was transformed with the pQE80L empty vector (-) or containing *appY* (WT) or mutant (K170E). Cultures were grown to OD_600_~0.5 and induced with 1 mM IPTG for 1 hour. RNA were extracted from 10^9^ cells and qRT-PCR experiments were performed. The genetic organization of flagellar genes is shown above the graph according to Ecocyc. Expression of the first gene of each operon was tested (*fliALF* in green, *flgB* in dark green, *flhD* in blue); *appC* (grey) is a control. Data are means +/− standard deviation (n=3). B. AppY overproduction affects motility. Overnight cultures were adjusted to OD_600_=1 and 1μl was spotted on plates without (light green) or with IPTG (green). Plates were incubated at 30°C during 15 to 20 h and swimming diameters were measured. A representative picture of swimming is shown below the graph. C. AppY favors FlhC degradation by the HslUV protease. MG1655 Δ*rpoS* (yellow) and MG1655 Δ*rpoS ΔhslU* (blue) strains containing *flhC-SPA* on the chromosome were transformed with the pQE80L empty vector (-, hatched) or containing *appY* (WT, plain) or mutant (K170E, dotted). Cultures were grown to an OD_600_~0.6 and induced with 1 mM IPTG for 1 hour. FlhC levels were analyzed by Western blotting using anti-Flag antibody. HtpG was detected as a loading control.

The global downregulation of a large number of genes involved in flagella synthesis suggests that FlhDC, the master regulator of this pathway, could itself be affected by AppY overproduction. However, neither the RNA-Seq nor the qRT-PCR experiments showed any significant decrease in *flhDC* expression when AppY was overproduced (Fig 5A, blue bars and Table S1). Therefore, the repression of the *fli* and *flg* operons cannot be attributed to a direct AppY effect on *flhDC* transcription or mRNA stability. Interestingly, FlhD and FlhC have been previously shown to be actively turned-over by the Lon and ClpXP proteases in *Proteus mirabilis* and *Salmonella enterica* Typhimurium, respectively (47,48). In order to follow the level of FlhDC in the presence of AppY, we fused a SPA-tag to FlhD and FlhC C-terminal extremities; for unknown reasons, this tag only led to FlhC detection. This absence of detection of FlhD has already been described (48). In the presence of the empty vector, we observed a constant level of FlhC with or without IPTG (Figure 5C). When AppY was overproduced, a decrease in FlhC amount was visualized; this decrease did not occur with the AppY _K170E_ mutant (Figure 5C). This result shows that AppY production favors FlhC degradation. Looking at our RNA-Seq experiments, we noticed that the expression of the *hslUV* protease was greatly enhanced in the presence of AppY (4.7 and 6-fold respectively). In order to determine if this protease was responsible for FlhC degradation, we looked at the FlhC level in strains deleted for *hslU*. This deletion led to FlhC-SPA stabilization even in the presence of AppY (Figure 5C). Overall, the data presented here suggest that AppY production results in the degradation of FlhC, which is at least partially dependent on HslUV. This active degradation leads to a severe motility defect in *E. coli*.

## DISCUSSION

In this paper, we have characterized AppY, a transcriptional regulator whose gene is carried by the DLP12 prophage. After identifying AppY direct and indirect targets, we have dissected the regulatory cascade leading to the modulation of three major processes, *i.e*. acid stress resistance, biofilm formation and motility. Our results, summarized in Figure 6, provide evidence for the first time for how, at the molecular level, a transcriptional regulator acquired by horizontal gene transfer integrates into the host regulatory network at several entry points to influence bacterial physiology.

**Figure 6 :**
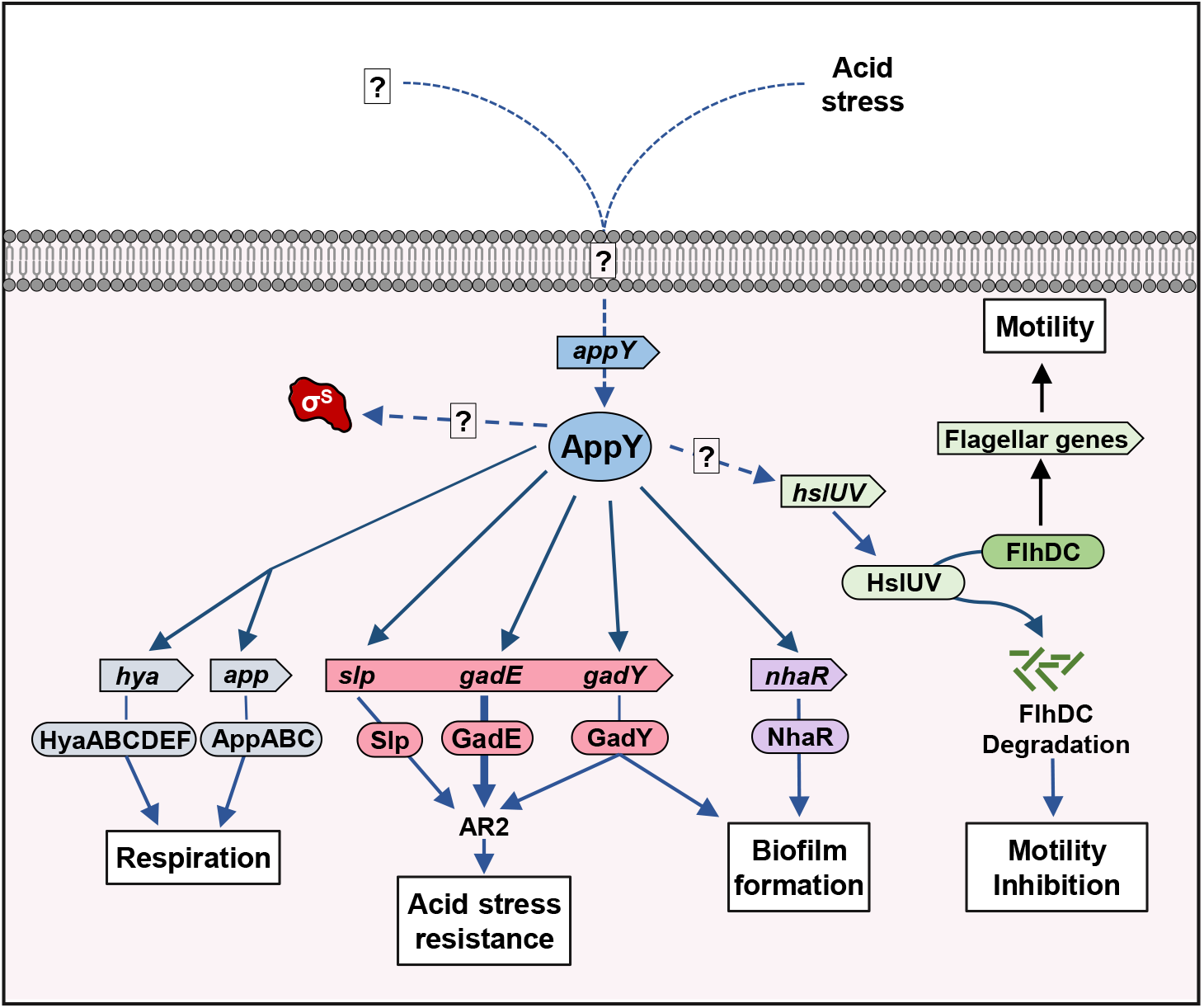
Pleiotropic effects of AppY on *E. coli* physiology. AppY activates several regulatory cascades leading to acid stress resistance, biofilm formation and motility repression. AppY also directly activates the expression of two operons involved in respiration but the consequences of these regulations on host physiology still have to be clarified. *appY* expression is induced by acid stress and probably by other environmental signals. AppY also leads to the stabilization of the sigma factor RpoS by an unknown mechanism. Under certain conditions, RpoS contributes to the expression of several genes under AppY control. The blue arrows indicate the regulations discovered in this work. The dotted arrows represent the unknown regulation.

A large number of the newly identified AppY-regulated genes are involved in the acid stress response (Figures 1–3 and 6). Pathogenic as well as commensal *E. coli* strains have developed a great resistance to acid stress, certainly to cope with the acidic environment of the stomach in order to reach the mammalian gut. Five distinct acid resistant systems, named AR1 to AR5, contribute to acid resistance in *E. coli* (49–51). These systems are not equally effective in dealing with acid stress; their efficacy varies depending on the medium composition and pH, as well as on the physiological state of the bacteria (33). The majority of AppY regulated genes are part of the acid fitness island and contribute to the AR2 pathway (Figures 1 and 2) (36,37,52). The regulation of this pathway is complex and involves a large number of regulators, including three transcriptional regulators from the AraC/XylS family: GadW, GadX and YdeO (33,36,53,54). In this study, we identify a fourth member of this family, namely AppY from prophage origin, which is also involved in the regulation of the AR2 pathway. The redundancy between the genes under the control of these 4 transcriptional regulators could explain why a strain deleted of *appY* still survived under acid stress conditions specific of the AR2 system (Figure S2C). Such functional redundancy has been previously shown for other AraC/XylS transcriptional regulators such as MarA, SoxS and Rob that can functionally substitute for one another (24).

The similarity between the AppY and the YdeO regulons is particularly striking and reinforces the idea that these two proteins could have similar functions (36,54). One major difference between these two regulators is that *ydeO* expression depends on the two-component system EvgS/EvgA, whereas *appY* expression does not rely on this system (Figure S2B) (55). This suggests that each of these regulators is involved in distinct responses triggered by specific signals. The EvgA-YdeO regulatory circuit is active in exponential phase, in a minimal medium containing glucose but is not responsible for AR2 induction in stationary phase or in rich medium, suggesting a possible role for AppY under these conditions (44). AppY has also been shown to be active under several other conditions such as carbon starvation, phosphate starvation or anaerobiosis (18,25). However, in our experimental conditions, *appY* was not induced under anaerobiosis as previously described; the reason for this discrepancy is not known (Figure S2A). In this study, we identified low pH as an additional signal triggering *appY* expression (Figure 3B). Performing RNA-Seq experiments under AppY-overproducing conditions was a necessary step to fully appraise the AppY regulon. It is now appealing to follow the dynamic of this regulon under the different environmental conditions leading to *appY* expression. This will allow us to determine if all AppY target genes are regulated under all conditions, if their expression varies depending on the encountered stress or if several simultaneous stresses are needed.

AppY is not the only transcriptional regulator from the AraC/XylS family from prophage origin that confers a selective advantage to its host under low pH. Indeed, in the enterohemorrhagic *E. coli* O157:H7 strain, two transcriptional regulators from this family located on a prophage, PsrB and PatE, also regulate genes from the AR2 system and enhance bacterial resistance to acidic conditions (56,57). In addition, PsrB and PatE also indirectly down-regulate the expression of genes coding for the type 3 secretion system, possibly in order to save energy (56–58). More generally, other transcriptional regulators acquired by horizontal gene transfer are involved in the regulation of ancestral genes. For example, some of them have been shown to antagonize the transcriptional silencing exerted by H-NS or, more recently, to activate the PhoP/PhoQ two component system, favoring here again the growth of their host under acidic pH (59,60)

In addition to its role in acid stress tolerance, our results also identify a role for AppY in biofilm formation (Figures 4, 6 and S3). Biofilm formation is a complex process in which many cellular components are involved and act depending on the environmental conditions (61). These cellular components include flagella, curli, exopolysaccharides and colonic acid. We have demonstrated that the increased biofilm formation observed when AppY is overproduced is due to the simultaneous and direct positive regulation of *nhaR* and *gadY* expression. *nhaR* codes for a transcriptional regulator belonging to the LysR family and *gadY* for a small regulatory RNA. These two factors have been shown to regulate the *pgaABCD* operon which is responsible for the synthesis and export of the exopolysaccharide ß-1.6-poly-N glucosamine (PGA) involved in biofilm formation (40,41,62). NhaR regulates this operon by directly binding to the *pgaABCD* promoter region, whereas GadY has been shown to titrate CsrA, a negative regulator of this operon. The concomitant activation of the AR2 system and of biofilm formation clearly shows that AppY activates different pathways to protect the cell from unfavorable environmental conditions.

AppY overproduction also caused a decrease in motility. Indeed, AppY overproduction massively represses the expression of genes coding for proteins involved in flagella synthesis. We demonstrated that this down-regulation is not due to the direct binding of AppY on the promoter regions of the flagellar genes but rather to an indirect effect on the master regulator of the flagella synthesis FlhDC (Figures 5 and 6). Our data showed that AppY overproduction increases the expression of the *hslUV* genes (4.7 and 6-fold respectively), which code for HslUV, one of the five energy-dependent proteases in *E. coli*, initially identified as a member of the heat shock regulon (63,64). HslUV increase in the cell leads to a partial FlhDC degradation (Figure 5C). However, according to the ChIP-Seq data, AppY does not directly regulate the *hslUV* operon (Figure 2A). This suggests that an additional factor regulated by AppY and controlling *hslUV* expression still remains to be identified. FlhDC degradation has already been observed in *Proteus mirabilis* and *Salmonella enterica* Typhimurium (47,48,65). In these organisms the degradation depends on the Lon and ClpXP proteases respectively. According to our data, AppY overproduction led to a mild increase in *lon* expression (2.5-fold) whereas *clpP* level remained constant in all tested conditions (Table S1). This suggests that in *E. coli*, ClpXP does not play any role in FlhDC degradation whereas Lon may still contribute (Table S1). This would be in accordance with the redundant role of HslUV and Lon already described for other substrates (64) The regulatory cascades leading to FlhDC degradation in *Proteus mirabilis* and *Salmonella enterica* Typhimurium have not been described; it would be interesting to investigate if a protein from prophage origin could be involved as well. Indeed, the regulation of the master regulator FlhDC by a protein from prophage origin is not an isolated example. For instance, the Sp5 prophage inhibits the motility of the *E. coli* MG1655 strain by repressing *flhDC* expression, but also by decreasing FlhDC amount in the cell. To date, the protein encoded by Sp5 and responsible for this regulation has not been identified (66). Motility regulation by morons is not restricted to swimming motility since twitching motility as well as swarming motility have all been described to be under the control of prophage genes (67–70) This down-regulation of the flagella synthesis is relevant with biofilm formation and could also be a way to save energy under unfavorable growth conditions. This fits, once again, with an overall involvement of AppY in bacterial survival.

Prior to this study, the *app* and *hya* operons were the only characterized AppY targets (18,19,25–28). Identifying these genes in the ChIP-Seq experiments both validates our approach and shows for the first time a direct AppY binding to the promoter regions of both operons (Figures 2B and 6). These two operons encode the cytochrome bd-II oxidase and the hydrogenase-I which are part of the respiratory machinery suggesting that AppY has also a direct role in regulating bacterial respiration. Interestingly, in addition to anaerobiosis, expression of the *hya* operon is also induced under acidic conditions (pH=5.5) and this is partially dependent on AppY (71). In another connection, a strain deleted for *cydB*, which codes for a subunit of the cytochrome bd-I oxidase, undergoes respiratory stress and displays a significant up-regulation of the *app* and *hya* operons as well as of the AR2 system and a down-regulation of flagellar genes (72). The major overlap between these transcriptomic data and the AppY regulon described here suggests a strong interaction between respiratory stress and acid stress that probably needs to be investigated in more detail to fully understand AppY contribution to bacterial physiology.

One striking observation from our ChIP-Seq data is that most of the AppY targets are also part of the RpoS regulon (73). This raises the question of the functional link between these two regulators. Further experiments are needed to determine if RpoS and AppY work together or regulate the same pool of genes under different physiological conditions. RpoS is a sigma subunit of the RNA-polymerase allowing the transcription of an important number of genes (29). When bacteria are growing in favorable conditions, RpoS is actively degraded by the ClpXP protease in the presence of the adaptor protein RssB (74–77). Under stressful conditions however, different members of a family of proteins called Ira block RpoS degradation by sequestering RssB. So far, three Ira proteins have been identified in *E. coli:* IraP, IraD and IraM (30,78). A previous study has shown that AppY also regulates this RpoS degradation pathway, independently of the known Ira proteins (30). Our data confirm that AppY production does not up-regulate the expression of *iraP*, *iraD* or *iraM* (Table S1). Since AppY regulates RpoS and RpoS has a huge impact on gene expression profiles, almost all the experiments presented here were performed in strains deleted of *rpoS*. Given the overlapping regulon between RpoS and AppY, these conditions were undoubtably necessary to unveil the full AppY regulon. The stabilization of RpoS along with induction of the AR2 system suggest that AppY can induce both a specific and a general stress response. Several questions remain concerning the chronology of these responses: is there a hierarchy, with first a specific response followed by the general stress response if the applied stress persists? Does the activation of one or the other depend on different “state” of AppY or of the encountered stress? Do they occur simultaneously?

The work presented here highlights the significant impact that a viral regulator can have on its bacterial host. However, we can predict that it represents only a small portion of the huge regulatory network existing between ancestral bacterial genes and genes from viral origin or, more widely, genes horizontally acquired. Consequently, studying the dialogue between phage and bacteria is key to fully understanding bacterial physiology and how it influences bacterial evolution, environment and human health.

## MATERIALS AND METHODS

### Media and Growth conditions

Cells were grown in Luria-Bertani (LB) broth. For acid stress experiments, we used E medium: (10 g/L MgSO_4_, 100 g/L citric acid, 500 g/L KH_2_PO_4_ and 175 g/L NaNH_4_HPO_4_ 4H_2_O), EG medium (E plus 0.4% D-Glucose) or potassium-modified LB (LBK; 10 g/L tryptone, 5 g/L of yeast extract, 7.45 g/L KCl)(79,80).

Liquid cultures were grown in aerobic conditions at 37°C under shaking (180 rpm) and plates were incubated at 37°C unless otherwise stated. The usual agar concentration for solid media is 1.5 %. When required, antibiotics were added at the following concentrations: 100 μg/mL ampicillin (Amp), 25 μg/mL chloramphenicol (Cam), 50 μg/mL kanamycin (Kan), 10 μg/mL tetracycline (Tet) and 25 μg/mL zeocin (Zeo).

### Bacterial strains, plasmids and primers

Bacterial strains are listed in Table S2, plasmids in Table S3 and primers in Table S4. All strains are derivatives of *E. coli* str. K-12 substr. MG1655, constructed by P1 transduction, selected on the appropriate antibiotic and verified by PCR. To construct the *flhC-SPA-kan* strain (where SPA is a Sequential Peptide Affinity tag) or the deletion/insertion mutants of *gadE, gadY* and *nhaR*, we used the *λ* red recombination system (81,82) (see more details in Supplemental Materials and Methods). The P-*appY* translational fusion was constructed using the NM580 strain, which contains the mini-*λ* prophage and has been engineered to construct chromosomal *lacZ* fusions using a selection/counter-selection method (83) (see more details in Supplemental Materials and Methods). This fusion contains a fragment ranging from −615 bp to +24 bp relative to the ATG start codon of *appY* Open Reading Frame (ORF). Site-specific mutagenesis was performed to introduce the K170E mutation into pQE-AppY (primers BAΦ184/BAΦ185; codon AAA changed to AGA) (84). Plasmids pND-671, pND-677 and pND-678 contain DNA fragments ranging from −598 pb to +14 pb relative to the *gadE* ATG start codon, from −245 bp to +14 bp relative to the beginning of the *gadY* sequence and from −210 bp to +15 bp relative to the ATG start codon for *gadA*. Plasmid pND-665 contains the full length *gadY* sequence and 245 nucleotides upstream. As we wanted to express *nhaR* from its own promoter region to keep the AppY-dependent induction and since it is in operon with *nhaA*, we amplified both genes plus a region of 482 nucleotides upstream *nhaA* ATG. PCR fragments were then digested with ClaI/BamHI for *gadY* and EcoRV/SalI for *nhaR* and cloned into pACYC184 digested with the same enzymes. The resulting plasmid is pND-692. All constructs were confirmed by Sanger sequencing.

### Chromatin Immunoprecipitation Sequencing (ChIP-Seq)

The ChIP-Seq protocol was adapted from (54,85). ND3 strain transformed with the pQE80L empty vector or containing untagged *appY, appY-3FLAG* or *appY*_K170E_-3FLAG, was grown in LB/Amp medium to an OD_600_ of 0.2. Cultures were then induced with 0.05 mM IPTG and further grown for 1 h. This experiment was performed with 3 independent cultures (replicates R1, R2 and R3) for each condition. Cells were fixed with 1 % formaldehyde; the crosslinking reaction was stopped by adding 250 mM glycine to the cultures. After being washed with PBS, the pellets were flash-frozen in liquid nitrogen and stored at −80°C. Cellular pellets were re-suspended in 1 mL of cold lysis buffer (50 mM Tris-HCL pH 7.5, 150 mM NaCl, 1 mM EDTA, 1% Triton-X100, protease inhibitor cocktail) plus 3.5 units of lysozyme (Novagen). Samples were then incubated at room temperature for 10 min on a rotating wheel (20 rpm) before shearing the crosslinked DNA by sonication using a Bioruptor Standard Waterbath Sonicator (Diagenode). After centrifugation, 900 μL of the supernatant were incubated with 40 μL of Agarose Anti-FLAG M2 gel beads (Sigma-Aldrich) and incubated overnight (ON) on a rotating wheel (20 rpm). After a gentle centrifugation step, the beads were washed sequentially with Low salt washing buffer, High salt washing buffer, LiCl washing buffer and twice with TBS buffer (see detailed composition in Supplemental Materials and Methods). Two elution steps of the FLAG-tagged protein-DNA complexes were performed with 200 μL of FLAG buffer (TBS buffer plus 100 μg/ml of 3XFLAG peptide from Sigma-Aldrich) and the samples were incubated 30 min at RT on a rotating wheel (20 rpm). After centrifugation, the protein-DNA crosslink was reversed by adding 25 μL of 5 M NaCl to the eluate and incubating ON at 65 °C. Proteins and RNA were removed by adding sequentially proteinase K and RNAse A (0.2 mg/ml). DNA was extracted with phenol/chloroform/isoamyl alcohol (25:24:1, Sigma-Aldrich) and ethanol precipitated. The DNA was finally resuspended in 30 μl of DNAse-free water, quantified with the Qubit^™^ dsDNA HS Assay kit (Invitrogen). DNA profiles were recorded with the TapeStation 4200 System (Agilent) in combination with the D5000 ScreenTape (Agilent). Libraries for high throughput DNA sequencing were prepared using the TruSeq^®^ChIP Sample Preparation kit (Illumina) according to the manufacturer’s protocol starting with 10 ng dsDNA from the previous step. High throughput DNA sequencing was performed as described below.

### RNA sequencing (RNA-Seq)

The ND3 strain transformed with pQE80L or the pQE80L-*appY* was grown in LB/Amp medium to OD_600_ = 0.6 at 37°C under shaking (180 rpm). 1 mM IPTG was then added and cells grown for one additional hour. Total RNAs were then purified as described above. This experiment was performed with 3 independent cultures (replicates R1, R2 and R3) for each condition. Total RNAs samples (3 μg) were depleted from ribosomal RNA using the bacteria Ribo-Zero^®^ rRNA Removal kit (Illumina) according to the manufacturer’s protocol. The cDNA libraries were then prepared from the depleted RNA samples obtained during the previous step using the GoScript^™^ Reverse transcriptase (Promega) to synthesize the first strand cDNA and the TruSeq^®^Stranded mRNA Library Prep kit (Illumina) according to the manufacturers’ protocols. High throughput DNA sequencing was performed as described below.

### RNA preparation and Reverse Transcription

For RNA-Seq and RT-qPCR experiments, cellular pellets (5.10^9^ cells) were flash-frozen in liquid nitrogen and stored at −80°C. The cells were harvested and frost at −80°C. Total RNAs were isolated from the pellet using the Maxwell^®^ 16 LEV miRNA Tissue Kit (Promega) according to the manufacturer’s instructions and an extra TURBO DNase (Invitrogen) digestion step to eliminate the contaminating DNA. The RNA quality was assessed by TapeStation 4200 system (Agilent). RNA was quantified spectrophotometrically at 260 nm (NanoDrop 1000; Thermo Fisher Scientific). For cDNA synthesis, 1 μg total RNA and 0.5 μg random primers (Promega) were used with the GoScript^™^ Reverse transcriptase (Promega) according to the manufacturer instruction.

### High throughput DNA sequencing

Prior to sequencing, RNA-Seq and ChIP-Seq libraries were quantified with the Qubit^™^ dsDNA HS Assay kit and their size distribution profiles recorded with the TapeStation 4200 System (Agilent) in combination with the D5000 DNA ScreenTape System (Agilent). Libraries were then diluted at 4 nM. Paired-end (2 × 75 bp) DNA sequencing was performed on the in-lab MiSeq sequencer hosted at the Institute Transcriptomic and Genomic facility with a MiSeq v3 (150-cycles) flow cell according to Illumina’s protocol. Table S5 summarizes the different sequencing runs performed during this study and the corresponding data yield for each sample. Raw sequencing reads (FASTQ files trimmed from their Illumina adaptors) were submitted to the NCBI Sequence Read Archive under the BioProject accession number PRJNA751735.

### High throughput sequencing data analysis

The FASTQ files generated by the MiSeq sequencer were trimmed and clipped for quality control with Trimmomatic (86) using the following parameters: ILLUMINACLIP:TruSeq3-SE:2:30:10 LEADING:3 TRAILING:3 SLIDINGWINDOW:4:15 MINLEN:36. For subsequent analyses we retained only the paired-reads FASTQ files generated by Trimmomatics.

### RNA-Seq analysis

Trimmed paired-reads were mapped on the *E. coli* MG1655 reference genome (NC_000913.3) and the raw read count per genomic feature computed with Rockhopper (87) using the following parameters: Orientation of mate-pair reads = rf, Max bases between paired reads = 500, Allowed mismatches = 0.15, Minimum seed length = 0.33, Min reads mapping to transcript = 10, Min transcript length = 50, Min count to seed a transcript = 50, Min count to extend a transcript = 5. Rockhopper mapping statistics are summarized in Table S6. We observed fairly high variations in the efficiency of the ribo-depletion with as much as 36% of the reads mapping to rRNA in one sample. In order to obtain a proper data normalization in downstream analyses, we manually cured the read count matrix generated by Rockhopper from remaining rRNA read counts (16S-, 23S- and 5S-rRNA); the resulting trimmed matrix was then use as input for the differential gene expression analysis performed with Voom/Limma method. The latter is embedded in DEGUST (https://degust.erc.monash.edu/), an interactive web-tool for RNA-Seq analysis (https://doi.org/10.5281/zenodo.3258932), that allows to compare 3 different methods for differential gene expression analysis as well as easy data visualization, browsing and export. All the RNA-Seq results are accessible on NCBI Sequence Read Archive under the BioProject accession number PRJNA751735. The RNA-Seq data are summarized in Table S1.

### ChIP-Seq analysis

Trimmed paired-end reads were mapped back onto the *E. coli* MG1655 reference genome (NC_000913.3) using Bowtie2 (88). The alignments were visualized and compared with IGV (89). Normalization and peak calling to identify enriched loci along the genome were done with the SeqMonk software (https://www.bioinformatics.babraham.ac.uk/projects/seqmonk/).

### Measure of expression with transcriptional *gfp* fusions

Strains ND3 or BAPHI089 were co-transformed with a plasmid carrying a *gfpmut2* transcriptional fusion and the pQE80L vector producing AppY wild-type (WT) or AppY_K170E_. ON cultures were diluted 60-fold into fresh LB medium supplemented with Amp, Kan and IPTG (0.05 mM) into a black 96-well plate with transparent bottom (Greiner). The plate was incubated into a Tecan Spark plate reader at 37 °C with shaking (180 rpm) during 10 h. Every 10 min, the optical density at 600 nm as well as the fluorescence intensity (excitation: 480 nm; emission: 535 nm) were measured. The transcriptional *gfp* fusion expression levels were standardized by dividing the fluorescence intensity by the OD600. Mean values and Standard deviation (SD) were computed on a minimum of three independent experiments.

### *appY* induction under acid conditions

This assay was carried out as previously described (42,43,79). Overnight cultures of the BAPHI018 strain carrying the *appY* translational fusion were diluted 1:1000 in LBK medium buffered with 100 mM piperazine-N, N’-bis-2-(ethanesulfonic acid) and adjusted at pH 5.7 or 8.5 using KOH.

Anaerobic cultures were performed in screw-caped test tubes with a slow agitation on a rotating wheel (8 rpm); aerobic cultures were performed in 14 ml aerated polypropylene tubes in an orbital water bath (200 rpm). At OD_600_ = 0.4, samples were taken for β-galactosidase assay using the method described by Miller (90). The pH of the cultures was verified at the end of the experiment to ensure that the values were maintained.

### Acid resistance assay

Acid resistance assays were performed as described previously (36). Briefly, overnight cultures were diluted 1:200 in 20 mL of LB at pH 7 with Amp and IPTG (1 mM) and grown at 37°C with aeration (180 rpm). At OD_600_ around one, 50 μL samples of culture were serially diluted in PBS and spotted on LB plates to count the colony forming cells (expressed in Colony Forming Unit or CFU). The resulting CFU represents the initial number of live cells before stress. In parallel, 50 μL of culture were transferred into 2 ml of preheated LB pH 2.5 (pH adjusted with HCl) and incubated for one hour at 37°C. 50 μL samples were serially diluted in PBS and 10 μL were spotted on LB plate. The CFU obtained here represents the cells having survived to the acid stress. The survival percentages were calculated by dividing the final CFU number by the initial CFU number. Each experiment was performed at least three times.

### Acid stress assay specific to the AR2 system

This assay was performed according to (35). Strains were grown for 22 hours at 37°C with aeration in LB medium with 0.4 % glucose at pH 7. These cultures were then diluted 1:1000 in 5 mL of EG medium at pH 2.2 (pH adjusted with HCL) supplemented with 1.5 mM sodium glutamate. As a control of the AR2 system induction, we used a medium corresponding to the EG medium at pH 2.2 without any additives. 100 μL samples of each culture were collected after 0, 2 and 4 hours of aerobic growth at 37°C. These samples were serially diluted in EG medium at pH 7, 10 μL were spotted on LB plate and incubated at 37°C.

### Quantitative Real-Time-PCR for Transcriptional analysis

Quantitative real-time PCR (qPCR) and the corresponding analysis were performed on a CFX96 Real-Time System (Bio-Rad). The reaction volume was 15 μL and the final concentration of each primer was 0.5 μM. The cycling parameters of the qRT-PCR were 98°C for 2 min, followed by 45 cycles of 98°C for 5 s, 55 °C for 10 s, 72°C for 1s. A final melting curve from 65°C to 95°C was performed to determine the specificity of the amplification. To determine the amplification kinetics of each product, the fluorescence derived from the incorporation of EvaGreen into the double-stranded PCR products was measured at the end of each cycle using the SsoFast EvaGreen Supermix 2X Kit (Bio-Rad, France). The results were analyzed using the Bio-Rad CFX Manager software, version 3.0 (Bio-Rad, France). The RNA16S gene was used as a reference for normalization. For each point a technical duplicate was performed. The amplification efficiencies for each primer pairs were comprised between 80 and 100%. All of the primer pairs used for qRT-PCR are reported in the Table S4.

### Motility assays

Motility assays were performed as described previously (36). Motility was tested on 0.5 % Tryptone and 0.5 % NaCl plates supplemented with 0.3% agar. Strains were grown overnight in LB with or without 0.5 mM IPTG. Overnight cultures were standardized to an OD_600_ = 1 and 1 μl of each culture were spotted on plates. Amp was added to the plates and to the cultures for plasmid maintenance when needed. Plates were incubated at 30 °C and after 15 to 20 h, the diameter of the swimming zone was measured using the ImageJ software (91). All strains were tested in triplicate and each experiment was independently performed three time.

### Biofilm quantification assay

Biofilm quantification was performed according to (92). Overnight cultures were diluted 1:1000 in fresh medium plus 0.5 mM IPTG in 24-well plate and incubated at 30°C without agitation for 24 h. Cultures were removed and the plate was washed two times with water before adding 0.1 % crystal violet (Sigma-Aldrich) for 10 min. The plate was washed three times with water, let dry overnight and samples were solubilized by using glacial acetic acid at 30 % (Carlo Erba). After an incubation of 15 min, the OD_550_ was measured to quantify biofilm formation. Standard deviations are based on a minimum of three independent experiments.

### Protein electrophoresis and Western blotting

Samples were analyzed using 12% lab-made standard SDS-polyacrylamide gels for AppY detection and using Precast Hepes-Tris gels 12% (WSHT) for other proteins. Semidry electrophoretic transfer into nitrocellulose membranes was performed using Trans-Blot Turbo Transfer System (BioRad). Membranes were probed with an anti-AppY antiserum (GenScript - 4.4:10000), or a Flag antibody (SIGMA - 1:10000) for FlhC-SPA detection and with HtpG antibody ((93)-1:100000) as a loading control. The blots were developed with Immobilon^®^Western (Millipore) using a chemiluminescence image analyzer (ImageQuant Las 4000).

## Acknowledgments

We thank all members of the Phage group in LCB, O. Genest, S. Gottesman and N. Majdalani for help and fruitful discussions. We are grateful to M.P. Castanié-Cornet for advice on acid stress experiments and G. Panis and O. Espeli for their help in setting up the ChIP-Seq experiments. We also thank N. O. Gomez for discussion on biofilm experiments and O. Genest for kindly providing the HtpG antibody. This work was supported by the Centre National de la Recherche Scientifique and Aix Marseille Université (AMU). A. B. was supported by a grant from the Agence Nationale de la Recherche (ANR-18-CE12-0024-01). N.D. is the recipient of a French Ministry fellowship.

