## Supplemental material for "Pleiotropic effects on *E. coli* physiology of the AraC-like regulator from prophage origin, AppY"

**SUPPORTING INFORMATION:**
**Supplemental Materials and Methods**

**Supplemental Figures:**
**Figure S1: Characterization of AppY-3FLAG WT and K170E**
**Figure S2: AppY contribution to acid stress**
**Figure S3: AppY favors biofilm formation *via* the direct induction of *nhaR* and *gadY*.**

**Supplemental Tables:**
**Table S1: Relative expression of genes under AppY overproduction (RNA-Seq data)**
**Table S2: Strains**
**Table S3: Plasmids**
**Table S4: Primers**
**Table S5: RNA-Seq and ChIP-Seq sequencing data**
**Table S6: Rockhopper mapping statistics**

### Supplemental Materials and Methods

#### Bacterial strains, plasmids and primers

To construct the *flhC*-SPA-*kan* strain or the deletion/insertion mutants of *gadY*, *nhaR* and *gadE* we used the λ Red recombination system. Briefly, the SPA-*kan*, the kanamycin resistance and the chloramphenicol resistance cassettes were amplified using as a template the BAPHI010 strain (primers BAΦ510/511), the BAPHI046 strain (primers BAΦ246/BAΦ247 for *gadY* and BAΦ359/BAΦ360 for *nhaR*) and the TKC strain (primers BAΦ084/BAΦ085) respectively (1–3). Primers used are 60 nucleotides long, they contain 20 nucleotides homologous to the desired cassette and 40 nucleotides homologous to the region flanking the coding sequences of *gadE*, *gadY* and *nhaR* for mutant's construction; for *flhC*-SPA-*kan* homology has to be on both sides of the STOP codon. The resulting PCR products were transformed into the NM1100 strain containing a mini-λ prophage and recombined into the bacterial chromosome thanks to the λ-red functions (4,5). Transformed cells were selected on the appropriate antibiotic and checked by PCR.
To construct the P-*appY* translational fusion, the primers used contains 40 nucleotides homologous to the region of insertion in NM580, corresponding to the zeocin cassette on the 5'end and to the *lacZ* ORF starting at the 9$^{th}$ codon on the 3' end. After transformation, the cells were recovered overnight on the bench in LB containing 1% glucose and plated on LB 1% arabinose to favor counter-selection. The resulting clones were screened for Kan$^{S}$, checked by PCR and sequenced. In the ND79 and ND85 strains, the gene conferring resistance to kanamycin was removed using pCP20 (6) to allow the transduction of other *kan* containing alleles.

#### Chromatin Immunoprecipitation Sequencing (ChIP-Seq)

Composition of buffers used for ChIP-Seq:
- Low salt washing buffer (0.1 % SDS, 1 % Triton-X100, 2 mM EDTA, 20 mM Tris-HCL pH 8.1, 150 mM NaCl)
- High salt washing buffer (0.1% SDS, 1% Triton-X100, 2 mM EDTA, 20 mM Tris-HCL pH 8.1, 500 mM NaCl)

- LiCl washing buffer (0.25 M LiCl, 1 % IGEPAL®, 1 % sodium deoxycholate, 1 mM EDTA, 10 mM Tris-HCl pH 8.1)
- TBS buffer (50 mM Tris-HCL pH 7.5, 150 mM NaCl).

Table S2: Strains

| Strain | Description | Reference or source |
|---|---|---|
| MG1655 | Wild-type *E. coli* | |
| BW25113 | *lacI+* $rrnB_{T14}$ $\Delta lacZ_{WJ16}$ *hsdR514* $\Delta araBAD_{AH33}$ $\Delta rhaBAD_{LD78}$ *rph-1 Δ(araB–D)567 Δ(rhaD-B)568 ΔlacZ4787(::rrnB-3) hsdR514 rph-1* | (1) |
| BA367 | MG1655 *crl- rpoS::tet* | Gottesman S., lab collection |
| C600 | *F-* $e14^-$ ($Mcr^-$) or $e14^-$ ($McrA^+$) *thr-1 leuB6 thi-1 lacY1 supE44 rfbD1 fhuA25* | (7) |
| NM580 | MG1655 *lacI-T1T2-*$zeo^R$*-pBRpLacO-kan-pBAD-ccdB. Mini-λ-Red::*$tet^R$ | (8) |
| NM1100 | *Mini-λ-Red::*$tet^R$ | (4) |
| SG22098 | MC4100 *clpP::cat* | (9) |
| TKC | *tet*, *kan*, *cat* containing *E. coli* strain | (2) |
| BAPHI010 | NM1100 AppY-SPA | NM1100 + PCR (BAΦ004/BAΦ005) |
| BAPHI018 | MG1655 $zeo^R$*-PappY*-SD-ATG + 8 codons *appY*-$9^{th}$ codon *lacZ* | NM580 + PCR (BAΦ001/BAΦ002) |
| BAPHI046 | MG1655 *lon::kan* | MG1655 + P1 (JW0429) |
| BAPHI056 | MG1655 *appY::kan* | MG1655 + P1 (JW0553) |
| BAPHI089 | MG1655 *rpos::tet gadE::cat* | ND3 + P1 (ND34) |
| ND2 | MG1655 *appY::kan rpoS::tet* | BAPHI056 + P1 (BA367) |
| ND3 | MG1655 *rpoS::tet* | MG1655 + P1 (BA367) |
| ND17 | MG1655 *gadE::cat* | MG1655 + P1 (ND34) |
| ND34 | NM1100 *gadE::cat* | NM1100 + PCR (BAΦ084/BAΦ085) |
| ND50 | MG1655 *gadY::kan* | NM1100 + PCR (BAΦ246/BAΦ247) |
| ND51 | MG1655 *rpoS::tet gadY::kan* | ND3 + P1 (ND50) |
| ND55 | MG1655 *nhaR::kan* | NM1100 + PCR (BAΦ359/BAΦ360) |
| ND56 | MG1655 *rpoS::tet nhaR::kan* | ND3 + P1 (ND55) |
| ND74 | MG1655 *rpoS::tet flhC-SPA-kan* | ND3 + P1 (*flhC*-SPA-*kan*) |
| ND76 | MG1655 *rpoS::tet clpP::cat flhC-SPA-kan* | ND78 + P1 (*flhC*-SPA-*kan*) |
| ND77 | MG1655 $zeo^R$*-PappY*-SD-ATG + 8 codons *appY*-$9^{th}$ codon *lacZ, evgA::kan* | BAPHI018 + P1 (JW2366) |
| ND78 | MG1655 *rpoS::tet clpP::cat.* | ND3 + P1 (SG22098) |
| ND79 | MG1655 *Δlon°* | BAPHI046 + pCP20 |
| ND80 | MG1655 *Δlon° rpoS::tet* | ND79 + P1 (BA367) |
| ND82 | MG1655 *Δlon° rpoS::tet flhC*-SPA° | ND80 + P1 (*flhC*-SPA-*kan*) |
| ND85 | MG1655 *rpoS::tet flhC*-SPA° | ND74 + pCP20 |
| ND86 | MG1655 *rpoS::tet hslU::kan flhC*-SPA° | ND85 + P1 (JW3902) |
| JW0429 | BW25113 *lon::kan* | (10) |

| | | |
|---|---|---|
| JW0553 | BW25113 *appY::kan* | (10) |
| JW1487 | BW25113 *gadC::kan* | (10) |
| JW2366 | BW25113 *evgA::kan* | (10) |
| JW3902 | BW25113 *hslU::kan* | (10) |

Table S3: Plasmids

| Plasmid | Description | Reference |
|---|---|---|
| | pQE80L, $Ap^R$, ColE1 replication origin, T5 promoter | Qiagen |
| | pQE80L-AppY, $Ap^R$ | (4) |
| pND-572 | pQE80L-$AppY_{K170E}$, $Ap^R$ | Directed mutagenesis on pQE80L-AppY with BAΦ184/BAΦ185 |
| pND-574 | pQE80L-AppY-3Flag, $Ap^R$ | PCR BAΦ075/BAΦ173, digested EcoRI/HindIII and inserted into pQE80L |
| pND-610 | pQE80L-$AppY_{K170E}$-3Flag, $Ap^R$ | Directed mutagenesis on pND574 with BAΦ184/BAΦ185 |
| | pUA66, $Km^R$, sc101 replication origin | (11) |
| | pUA66-PappC, $Km^R$ | (11) |
| | pUA66-P*gadB*, $Km^R$ | (11) |
| | pUA66-P*gadX*, $Km^R$ | (11) |
| | pUA66-P*hdeA*, $Km^R$ | (11) |
| | pUA66-P*hdeD*, $Km^R$ | (11) |
| | pUA66-P*hyaA*, $Km^R$ | (11) |
| | pUA66-P*slp*, $Km^R$ | (11) |
| pND-671 | pUA66-P*gadE*, $Km^R$ | PCR with BAΦ227/BAΦ228, digested XhoI/BamHI inserted into pUA66 |
| pND-677 | pUA66-P*gadY*, $Km^R$ | PCR with BAΦ236/BAΦ237, digested XhoI/BamHI inserted into pUA66 |
| pND-678 | pUA66-P*gadA*, $Km^R$ | PCR with BAΦ238/BAΦ239, digested XhoI/BamHI inserted into pUA66 |
| | pACYC184, $Cm^R$, p15A replication origin | Biolabs |
| pND-665 | pACYC184-*gadY*, $Cm^R$ | PCR with BAΦ371/BAΦ372, digested ClaI/BamHI and introduced into pACYC184 |
| pND-692 | pACYC184-*nhaA*-*nhaR*, $Cm^R$ | PCR with BAΦ584/BAΦ586, digested EcoRV/SalI and introduced into pACYC184 |

Table S4: Primers

| Primer number | Primer name | Primer sequence (5'-3') |
|---|---|---|
| BAΦ001 | Chromosomal *appY* translational *lacZ* fusion Fwd | TAC-TAT-GCC-GAT-ATA-CTA-TGC-CGA-TGA-TTA-ATT-GTC-AAC-GTA-TCG-GGT-GCT-GCT-AAA-CC |
| BAΦ002 | Chromosomal *appY* translational *lacZ* fusion Rev | CCA-GGG-TTT-TCC-CAG-TCA-CGA-CGT-TGT-AAA-ACG-ACG-GCA-ACT-ACG-GAG-CAA-ACA-TAA-TC |
| BAΦ004 | Chromosomal *appY*-SPA Fwd | AAA-ATA-ATC-GGC-GTC-ACA-GAT-GGA-ATA-AAC-AAA-ACA-ATT-GAC-TCC-ATG-GAA-AAG-AGA-A |
| BAΦ005 | Chromosomal *appY*-SPA Rev | TAT-TAT-AAT-TAA-CAT-GTA-GAC-AAC-TTG-TAA-TAA-ACA-TTA-CAT-ATG-AAT-ATC-CTC-CTT-AG |
| BAΦ075 | *appY*-3Flag Fwd | ATC-GGA-ATT-CAT-TAA-AGA-GGA-GAA-ATT-AAC-TAT-GGA-TTA-TGT-TTG-CTC-CGT-AGT-TTT-CAT |
| BAΦ084 | *gadE::cam* Fwd | GGA-TGA-CAT-ATT-CGA-AAC-GAT-AAC-GGC-TAA-GGA-GCA-AGT-TTG-TGA-CGG-AAG-ATC-ACT-TCG |
| BAΦ085 | *gadE::cam* Rev | CTC-GTC-ATG-CCA-GCC-ATC-AAT-TTC-AGT-TGC-TTA-TGT-CCT-GAA-CCA-GCA-ATA-GAC-ATA-AGC-G |
| BAΦ147 | qRT-PCR, *appC* Fwd | GAT-GGA-AGG-GGA-GTG-GCA-AA |
| BAΦ148 | qRT-PCR, *appC* Rev | CGT-GGG-TAG-GTT-TCA-GCC-AT |
| BAΦ149 | qRT-PCR, *fliA* Fwd | AGC-GAG-AAA-ACC-CGC-TAC-AA |
| BAΦ150 | qRT-PCR, *fliA* Rev | TGT-GTA-ACT-GAC-TGA-CCC-GC |
| BAΦ151 | qRT-PCR, *flgB* Fwd | CAA-TGC-CGA-TAC-CCC-TGG-TT |
| BAΦ152 | qRT-PCR, *flgB* Rev | TTG-CAG-TTC-TGC-GGT-AGG-AG |
| BAΦ153 | qRT-PCR, *fliF* Fwd | CTG-CGT-GCG-AAT-CCG-AAA-AT |
| BAΦ154 | qRT-PCR, *fliF* Rev | CTT-CGC-TGA-AGC-GGT-AAG-GA |
| BAΦ155 | qRT-PCR, *fliL* Fwd | TTT-GCC-GGA-AGT-CCG-TAG-TC |
| BAΦ156 | qRT-PCR, *fliL* Rev | TCG-GTG-ACA-TCC-TGT-TTC-GG |
| BAΦ173 | *appY*-3Flag Rev | GAT-CAA-GCT-TCT-ACT-TGT-CAT-CGT-CAT-CCT-TGT-AGT-CGA-TGT-CAT-GAT-CTT-TAT-AAT-CAC- |

| | | |
|---|---|---|
| | | CGT-CAT-GGT-CTT-TGT-AGT-CGT-CAA-TTG-TTT-TGT-TTA-TTC |
| BAΦ184 | AppY K170E Fwd | GAA-AGT-TTA-ATA-GAA-AAA-AGA-TTA-A |
| BAΦ185 | AppY K170E Rev | TTA-ATC-TTT-TTT-CTA-TTA-AAC-TTT-C |
| BAΦ227 | pUA66-*gadE* Fwd | GAT-GCT-CGA-GTT-ACC-CCG-GTT-GTC-ACC-CGG |
| BAΦ228 | pUA66-*gadE* Rev | GAT-CGG-ATC-CGT-CAT-GAG-AAA-AAT-CAT-AAC |
| BAΦ236 | pUA66-*gadY* Fwd | GAT-GCT-CGA-GGA-TTA-TCC-CTT-ATA-TTT-CAT-AC |
| BAΦ237 | pUA66-*gadY* Rev | GAT-CGG-ATC-CAA-CTT-TGT-GCT-CTC-AGT-AAG |
| BAΦ238 | pUA66-*gadA* Fwd | GAT-GCT-CGA-GTT-AAT-TTG-ATC-GCC-CGA-ACA-G |
| BAΦ239 | pUA66-*gadA* Rev | GAT-CGG-ATC-CAA-CAG-CTT-CTG-GTC-CAT-TTC |
| BAΦ246 | *gadY::k*an Fwd | AAT-GGC-TGA-TCT-TAT-TTC-CAG-TAA-AAG-TTA-TAT-TTA-ACT-TAT-TCC-GGG-GAT-CCG-TCG-ACC |
| BAΦ247 | *gadY::k*an Rev | CTG-CGG-AAG-GAA-TAA-GAT-TAT-AGA-GTT-TTA-CTC-AGA-CAT-ATG-TAG-GCT-GGA-GCT-GCT-TCG |
| BAΦ359 | *nhaR::k*an Fwd | GCC-ATA-AAC-GGC-TCC-CTT-TTC-ATT-GTT-ATC-AGG-GAG-AGA-AAT-TCC-GGG-GAT-CCG-TCG-ACC |
| BAΦ360 | *nhaR::k*an Rev | CGC-ACC-GCT-GGA-CTA-AAA-AGC-GCA-GAA-TAA-TCC-GTA-TTG-CTG-TAG-GCT-GGA-GCT-GCT-TCG |
| BAΦ371 | pACYC184-*gadY* Fwd | CTA-GAT-CGA-TGA-TTA-TCC-CTT-ATA-TTT-CAT-AC |
| BAΦ372 | pACYC184-*gadY R*ev | GAT-CGG-ATC-CAA-AAA-AAC-CCG-GCA-TAG-GGG |
| BAΦ376 | qRT-PCR, *flhD* Fwd | TCC-GCT-ATG-TTT-CGT-CTC-GG |
| BAΦ377 | qRT-PCR, *flhD* Rev | ATC-GTC-AAC-GCG-GGA-ATC-TT |
| BAΦ510 | Chromosomal *flhC*-SPA Fwd | TAT-CCC-ACA-ACT-GCT-GGA-TGA-ACA-GAG-AGT-ACA-GGC-TGT-TTC-CAT-GGA-AAA-GAG-AAG |
| BAΦ511 | Chromosomal *flhC*-SPA Rev | GTC-GTT-ACC-GCT-GCT-GGA-ATG-TTG-CGC-CTC-ACC-GTA-TCA-GCA-TAT-GAA-TAT-CCT-CCT-TAG |
| BAΦ584 | pACYC-*nhaA*-*nhaR* Fwd | CTA-GGA-TAT-CCT-ATC-TGC-CGT-TCA-GCT-AAT-G |
| BAΦ586 | pACYC-*nhaA*-*nhaR* Rev | GAT-CGT-CGA-CTT-AAC-GCA-CCG-CTG-GAC-TAA-AAA-G |

| | | |
|---|---|---|
| 16S-EC1 | qRT-PCR, 16S RNA Fwd | GTT-AAT-ACC-TTT-GCT-CAT-TGA |
| 16S-EC2 | qRT-PCR, 16S RNA Rev | ACC-AGG-GTA-TCT-AAT-CCT-GTT |

| Sequencing | Sample | Replicate | Run # | Data yield |
|---|---|---|---|---|
| RNA-Seq | ND3-pQE80L | R1 | 1 | 702.8 Mbp |
| | | R2 | 2 | 792.12 Mbp |
| | | R3 | 3 | 917.01 Mbp |
| | ND3-pQE80L-*appY* | R1 | 1 | 723.02 Mbp |
| | | R2 | 2 | 774.54 Mbp |
| | | R3 | 3 | 942.25 Mbp |
| ChIP-Seq | ND3-pQE80L | R0* | 4 | 306.40 Mbp |
| | ND3-pQE80L-*appY* | R0* | 4 | |
| | | R1 | 5 | |
| | | R2 | 6 | |
| | | R3 | 6 | |
| | ND3-pQE80L-*appY-3FLAG* | R0* | 4 | |
| | | R1 | 5 | |
| | | R2 | 6 | |
| | | R3 | 6 | |
| | ND3-pQE80L-*appY*$_{K170E}$*-3FLAG* | R1 | 5 | |
| | | R2 | 6 | |
| | | R3 | 6 | |

**Table S5: Sequencing runs performed during this study.** *Samples tested to assess the entire ChIP-seq workflow. These reads were neither included in the downstream data analysis nor submitted to NCBI.

| Construct | pQE80L | | | pQE80L-*appY* | | |
|---|---|---|---|---|---|---|
| **Replicate** | R1 | R2 | R3 | R1 | R2 | R3 |
| **Total reads** | 4541549 | 5124029 | 5914850 | 4637874 | 5010046 | 6094302 |
| **Successfully aligned reads** | 86% | 92% | 91% | 98% | 98% | 98% |
| **Aligning (sense) to protein-coding genes** | 54% | 83% | 76% | 88% | 86% | 86% |
| **Aligning (antisense) to protein-coding genes** | 1% | 1% | 1% | 1% | 1% | 1% |
| **Aligning (sense) to rRNA** | 36% | 3% | 13% | 2% | 2% | 2% |
| **Aligning (antisense) to rRNA** | 4% | 0% | 0% | 0% | 0% | 0% |
| **Aligning (sense) to tRNA** | 0% | 1% | 0% | 0% | 0% | 0% |
| **Aligning (antisense) to tRNA** | 0% | 0% | 0% | 0% | 0% | 0% |
| **Aligning (sense) to miscellaneous RNA** | 2% | 5% | 3% | 3% | 4% | 4% |
| **Aligning (antisense) to miscellaneous RNA** | 0% | 0% | 0% | 0% | 0% | 0% |
| **Aligning to unannotated region** | 4% | 7% | 6% | 5% | 6% | 6% |

**Table S6: Rockhopper mapping statistics.**

Supplemental Figures S1

A.

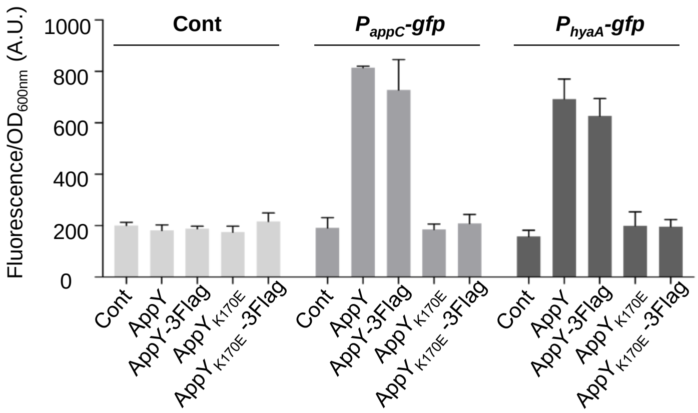

B.

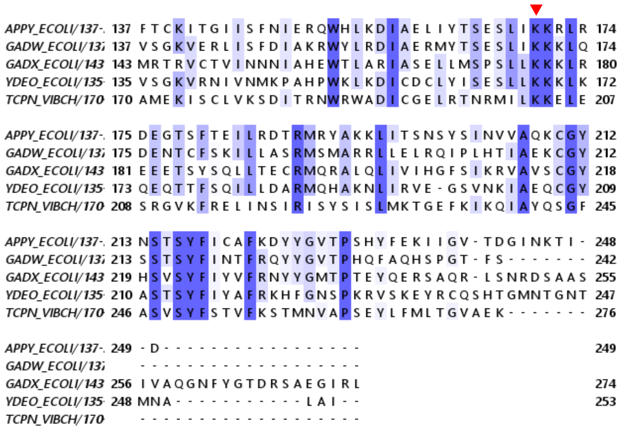

C.

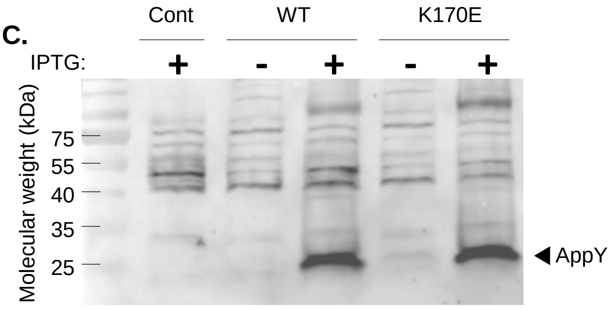

**Figure S1 : Characterization of AppY-3FLAG WT and K170E**

A. Functionality of the different AppY constructs used in this study. pQE80L and pQE-*appY*$_{WT}$ or pQE-*appY*$_{K170E}$, with or without the 3-Flag tag were co-transformed in MG1655 Δ*rpoS* with the pUA66 empty vector (Cont) or the transcriptional fusions P$_{appC}$-*gfp* and P$_{hyaA}$-*gfp*. Cells were grown in LB at 37°C with 0.05 mM IPTG during 10 hours. Activity of the fusions was determined as described in Materials and Methods. The mean of 3 replicates is presented here and the standard deviation (SD) is indicated by the error bars. A.U., arbitrary units. B. Alignment of AppY C-terminal domain with other proteins from the AraC family. Sequence alignment was made using Jalview (12). The intensity of the blue color reflects the residue conservation. The K170 residue mutated in this study is indicated by a red arrow. C. AppY$_{WT}$ and AppY$_{K170E}$ production. pQE80L empty vector (Cont) or containing *appY* (WT) or mutant (K170E) were transformed in MG1655 Δ*rpoS*. Cells were grown at 37°C in LB to OD$_{600}$ ≃ 0.6 and 0.05 mM IPTG was added for 1 hour. AppY levels were analyzed by Western blotting using an anti-AppY antiserum.

Supplemental Figures S2

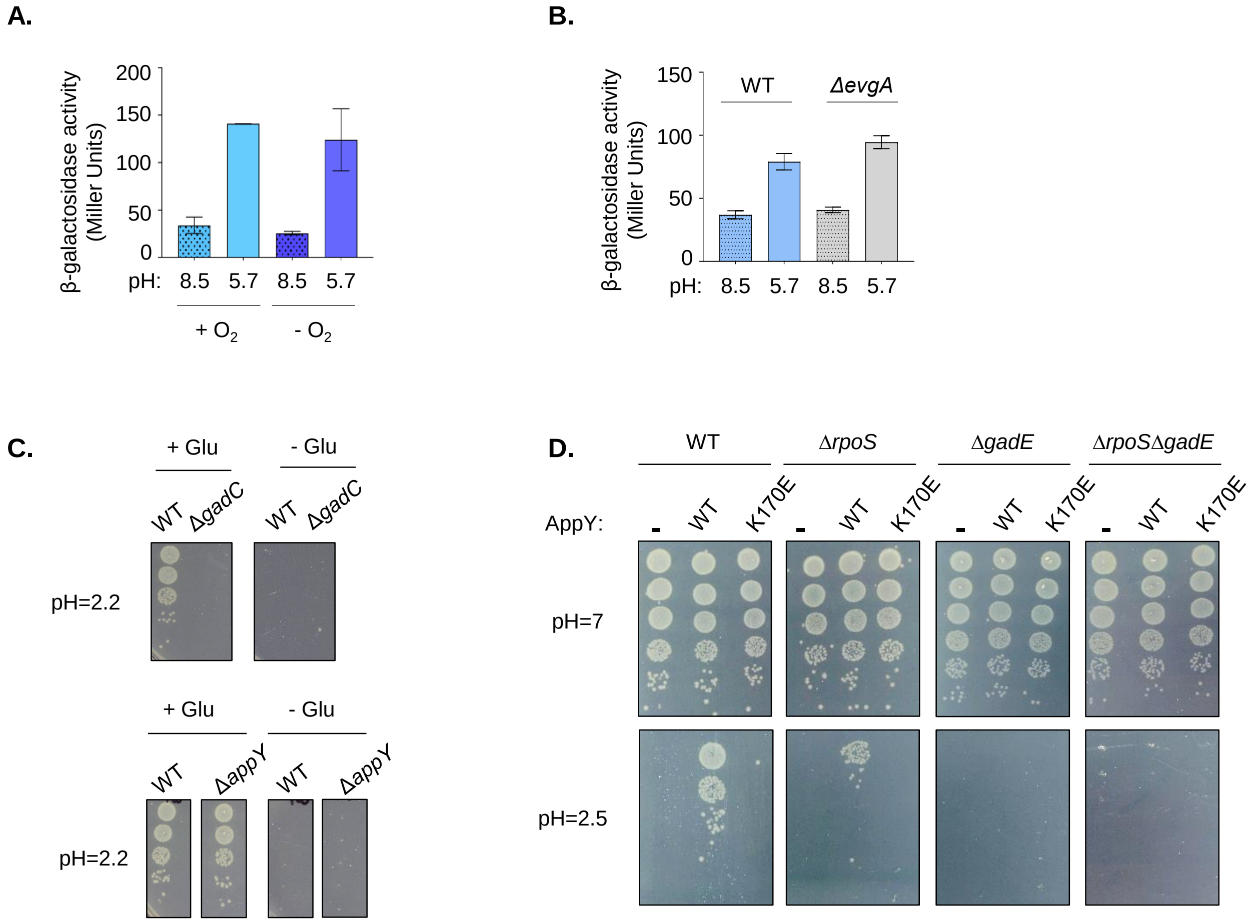

**Figure S2 : AppY contribution to acid stress**

A. Strains carrying a chromosomal *appY-lacZ* translational fusion were cultured overnight in LBK pH= 7, diluted 1:1000 into LB pH=8.5 (dotted line) or 5.7 (plain) and incubated at 37 °C in aerobic (light blue) or anaerobic (dark blue) conditions. Cultures were grown until an $OD_{600}$ ~ 0.4 . The activity of the *appY* fusion was determined as described using the Miller assay (13). Data are means +/- standard deviation (n=3). B. Experiments were performed as described in A with a WT (blue) and Δ*evgA* (grey) strains in aerobic conditions C. Strains BW25113 (WT) and Δ*gadC* or MG1655 (WT) and Δ*appY*, were grown in LB plus 0.4 % glucose at 37 °C for 22 hours. Cultures were diluted 1:1000 into EG medium pH= 2.2 and grown with or without sodium glutamate for 4 hours. Cells were serially diluted and 10 *µ*l of cultures were spotted on LB plate incubated at 37 °C. D. AppY overproduction confers resistance to acid stress. MG1655 (WT), Δ*rpoS* , Δ*gadE* or Δ*gadE* Δ*rpoS* strains transformed with pQE80L (-), pQE-$appY_{WT}$ (WT) or pQE-$appY_{K170E}$ (K170E) were grown to $OD_{600}$=1 in LB broth (pH 7.0) with 1 mM IPTG. Cells were diluted 40-fold into LB broth (pH 2.5) and incubated for 1 h at 37 °C. Cells were serially diluted and 10 µl of cultures were spotted on LB plate incubated at 37 °C.

Supplemental Figure S3

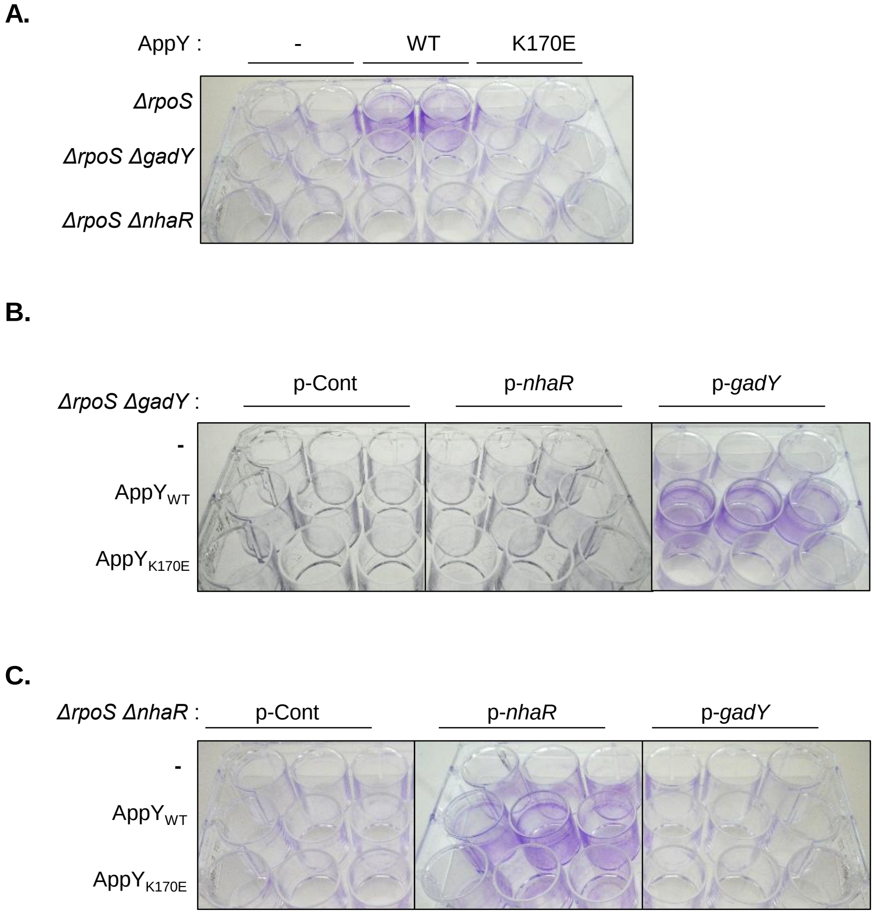

**Figure S3 : AppY favors biofilm formation *via* the direct induction of *nhaR* and *gadY*.**
The indicated strains were grown in LB plus 0.5 mM IPTG at 30°C without shaking for 24 hours. Biofilm was visualized using crystal violet staining. A. Biofilm formation dependent on NhaR and GadY. MG1655 Δ*rpoS*, MG1655 Δ*rpoS* Δ*gadY* or MG1655 Δ*rpoS* Δ*nhaR* were transformed with the pQE80L empty vector (-) or containing *appY* (WT) or mutant (K170E). B. Complementation of MG1655 Δ*rpoS* Δ*gadY* with *gadY and nhaR* expressed under their own promoter. MG1655 Δ*rpoS* Δ*gadY* strain was co-transformed with a pQE80L construct (empty vector (-), containing *appY* (WT) or mutant (K170E)) and a pACYC184 construct (empty vector (p-Cont), containing *nhaR* (p-*nhaR*) or *gadY* (p-*gadY*)). C. Complementation of MG1655 Δ*rpoS* Δ*nhaR* with *gadY and nhaR* expressed under their own promoter. MG1655 Δ*rpoS* Δ*nhaR* strain was co-transformed with a pQE80L construct (empty vector (-), containing *appY* (WT) or mutant (K170E)) and a pACYC184 construct (empty vector (p-Cont), containing *nhaR* (p-*nhaR*) or *gadY* (p-*gadY*)).
